# Cell-substrate distance fluctuations of confluent cells enable fast and coherent collective migration

**DOI:** 10.1101/2024.03.07.583903

**Authors:** Marcel Jipp, Bente D. Wagner, Lisa Egbringhoff, Andreas Teichmann, Angela Rübeling, Paul Nieschwitz, Alf Honigmann, Alexey Chizhik, Tabea A. Oswald, Andreas Janshoff

**Affiliations:** University of Göttingen, Institute of Physical Chemistry, Tammannstr. 6, 37077 Göttingen, Germany; University of Göttingen, Institute of Organic and Biomolecular Chemistry, Tammannstr. 2, 37077 Göttingen, Germany; Biotechnology Center, Technische Universität Dresden Pfotenhauerstraße 108, 01307 Dresden, Germany; University of Göttingen, Third Institute of Physics Friedrich-Hund-Platz 1, 37077 Göttingen, Germany

**Keywords:** tight junctions, traction force microscopy, collective cell migration, jamming, cell mechanics, atomic force microscopy

## Abstract

Collective cell migration is an emergent phenomenon, with long-range cell-cell communication influenced by various factors, including transmission of forces, viscoelasticity of individual cells, substrate interactions, and mechanotransduction. We investigate how alterations in cell-substrate distance fluctuations, cell-substrate adhesion, and traction forces impact the average velocity and temporal-spatial correlation of confluent monolayers formed either by wild-type MDCKII cells or zonula occludens (ZO) 1/2-depleted MDCKII cells (dKD) representing highly contractile cells.

The data indicates that confluent dKD monolayers exhibit decreased average velocity compared to less contractile WT cells concomitant with increased substrate adhesion, reduced traction forces, a more compact shape, diminished cell-cell interactions, and reduced cell-substrate distance fluctuations. Depletion of basal actin and myosin further supports the notion that short-range cell-substrate interactions, particularly fluctuations driven by basal actomyosin, significantly influence the migration speed of the monolayer on a larger length scale.

## Introduction

Biological tissues emerge from long-range collective morphological transitions during development and regeneration. Diverse manifestations of collective cell migration exist, contingent upon the specific biological tissue and process.^1–4^ In epithelial morphogenesis, wound healing, and regeneration processes, cells typically migrate in cohesive sheets attached to the extracellular matrix (ECM).^5,6^ Perturbations of cellular motion lead to severe developmental defects, compromised wound healing, or even tumor formation.^1,7,8^ Collective migration is a function of cell shape, cell-cell connectivity, cell-substrate adhesion, and active traction forces.^9,10^

Although the significance of adherens junctions (AJs) in creating contractile belts that connect cells into a seamless sheet and focal contacts formed between the cell’s basal side and the extracellular matrix has been extensively demonstrated,^11–14^ the importance of tight junctions (TJs) in the context of collective cell migration has only recently come under scrutiny. Tight junctions constitute multiprotein junctional complexes that create a seal between epithelial cells. Among the various proteins, peripheral membrane protein zona occludens-1 (ZO-1) is crucial for actomyosin organization and contractility.^15^ ZO-1 knockout (KO) cell lines exhibit an elevation in overall traction forces at the AJ due to the negative regulation of mechanical forces in the cells, leading to stronger cell-cell and cell-substrate forces.^16,17^ Depletion of both ZO-1 and ZO-2 (double knock-down, dKD) results in an even more noticeable phenotype. MDCKII dKD cells reveal a dramatically thickened peri-junctional actin ring leading to an increased apicolateral tension directly at the cell junctions.^18–21^ In contrast, apical surface tension in cell centers is decreased as excess surface area accumulates.^22,23^

Studies on the impact of functional tight junctions on cellular motility in migration have yielded conflicting evidence. KO studies targeting single ZO proteins have shown accelerated migration (ZO-1 in MCF-10A cells^17^; ZO-2 in MDCKII cells^24^) and impaired migration (ZO-1 in adult HDMEC-c cells^25^; ZO-3 in MCF-10A cells^17^). However, cellular migration is severely compromised when MDCKII cells lack the transmembrane TJ protein occludin^18^ or a combination of ZO-1 and ZO-2 in MDCKII cells^22^. In both cases, it could be shown that the force balance is disturbed. Further studies showed that the balance between cellular contractility and differential adhesion between cells leads to a time-dependent demixing in MDCKII cell (wild-type) WT and dKD confluent co-cultures.^26^ These findings show some discrepancies not only because of the different cell types used and the proteins chosen for tight junction interference, but also the differences in migration assays (scratch-assays, patterning, or stencils) used.

In this study, our objective is to delve into the effects of individual cell contractility and the changes in cell-substrate interactions on the collective migration of MDCKII monolayers. We show that ZO-1/ZO-2 dKD cells organized in confluent monolayers are generally slower and display less coordinated movement than WT cells, even in a pre-jammed state. We attribute the loss of collective cell rearrangement to reduced actively driven cell-substrate distance fluctuations. This is accompanied and fostered by a reduction in the traction forces required to surpass the enhanced adhesive strength observed in cells lacking ZO-1/ZO-2. Our findings emphasize the requirement of sufficient cell-substrate dynamics to enable large-scale, coherently fluid movement of cells in mesoscale clusters, which could also have implications for cellular migration in a wound healing scenario.

## Results

### WT cells show higher average velocities than dKD cells until the onset of jamming

The impact of compromised tight junctions and altered cell-substrate interactions on the average velocity of MDCKII monolayers was assessed by phase contrast microscopy and particle image velocimetry (PIV).^27^ Cells were cultivated on collagen-coated polyacrylamide gels (PAA), with a Young’s modulus of 1.61±0.01 kPa measured by atomic force microscopy (AFM). The gel substrate not only mimics the stiffness of native environments for cells but also enables us to quantify traction forces exerted by individual cells on the deformable substrate. Phase contrast microscopy images of confluent WT cells reveal a higher average velocity compared to dKD monolayers (SI Videos and Figure 1A). The average velocity was obtained from averaging the magnitude of velocity vectors from PIV analysis.

**Figure 1.**
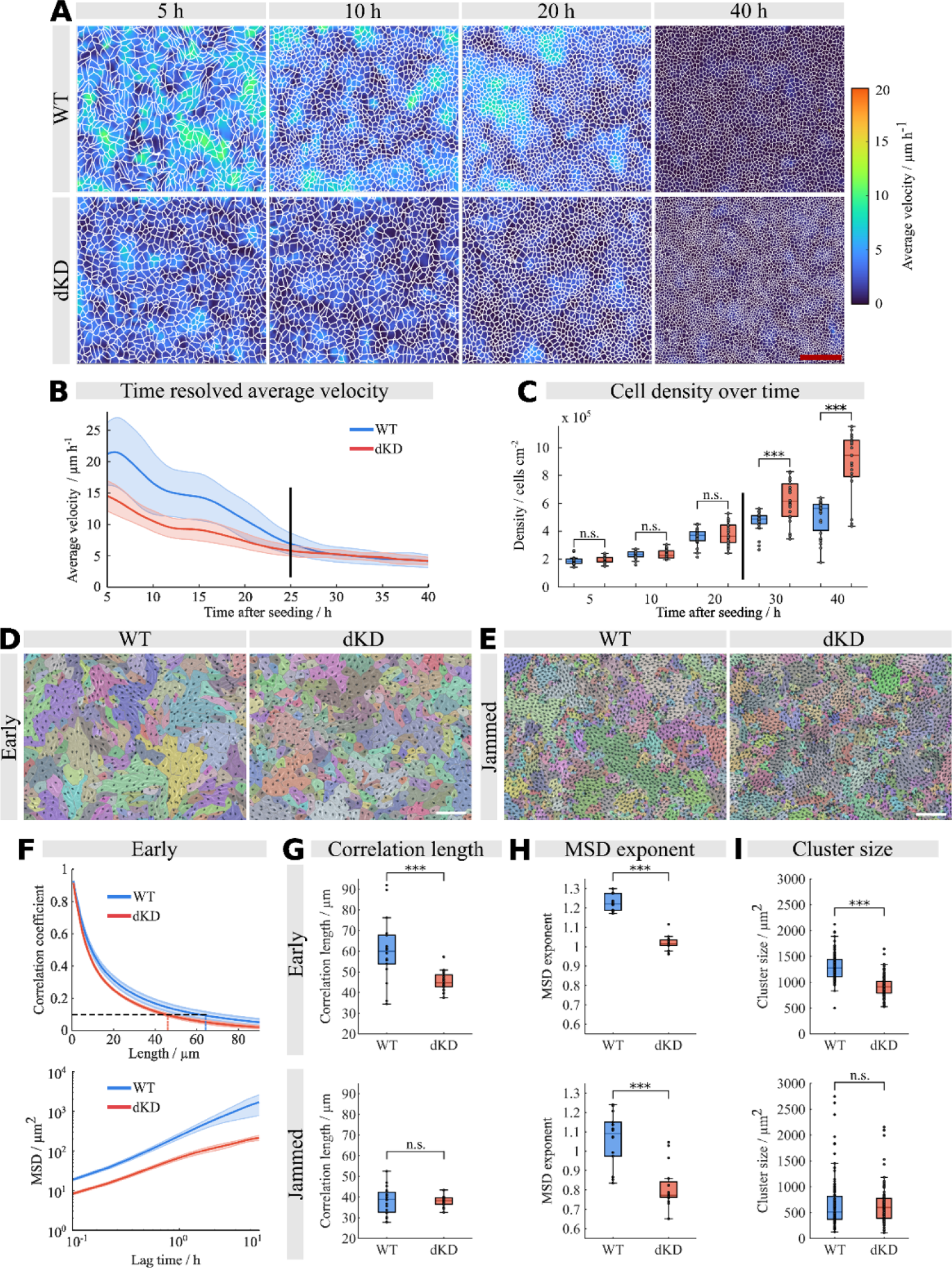
Motility is altered in tight junction deficient epithelia. A) Representative velocity maps generated from the corresponding velocity vectors obtained from PIV with cell outlines (white) obtained from Cellpose starting from the point of confluence (5 h) until 40 h after the experiment started. Scale bar: 160 µm. B) Corresponding averaged time-resolved velocities showing a “baseline velocity” being reached for all samples after approx. 30 h (black line). C) Cell densities (cells cm^−2^) obtained from Cellpose. The black line between 20 h and 30 h indicates the detection limit. D) Representative motion cluster maps of WT and dKD cells with corresponding cell outlines obtained by Cellpose analysis at early (5 h - 10 h after seeding) and jammed (30 h - 35 h after seeding) stages of the observation period. The arrows represent the motion of individual cells. Cells with similar directionality are grouped into a cluster with the same color. Scale bar: 150 µm. F) Upper plot: spatial correlation of velocity for early times. Vertical dashed lines indicate the corresponding correlation lengths, which are shown in G). Lower plot: Mean square displacements (MSD) of WT and dKD at early times. The corresponding MSD exponents are shown in H). I) Cluster sizes of early (upper plot) and jammed (lower plot) states. Each point represents the average cluster sizes of a single frame.

We examined intervals spanning all time frames (7.5 min/frame) commencing from the point at which confluency is reached (five hours after seeding). Overall, WT monolayers show a significantly higher average velocity of 21±5 µm/h compared to dKD cells (15±2 µm/h) at the point of confluence (Figure 1B). Throughout the observation period, the average velocity of both cell lines drops to a “baseline velocity” (WT: 4.5±1.0 µm/h, dKD: 4.7±0.7 µm/h at 35 h after seeding). This decay can be explained by contact inhibition,^28^ resulting in a gradual rise in cell density accompanied by a decrease in eccentricity (SI-Figure 1A-D), ultimately culminating in jamming.^29^ At this time (baseline velocity), the average cellular velocity drops below the resolution level of the PIV method, determined by assessing the average velocity of fixed, non-migrating confluent cells that also showed an average velocity of 4.2±1.1 µm/h (SI-Figure 2A).

**Figure 2.**
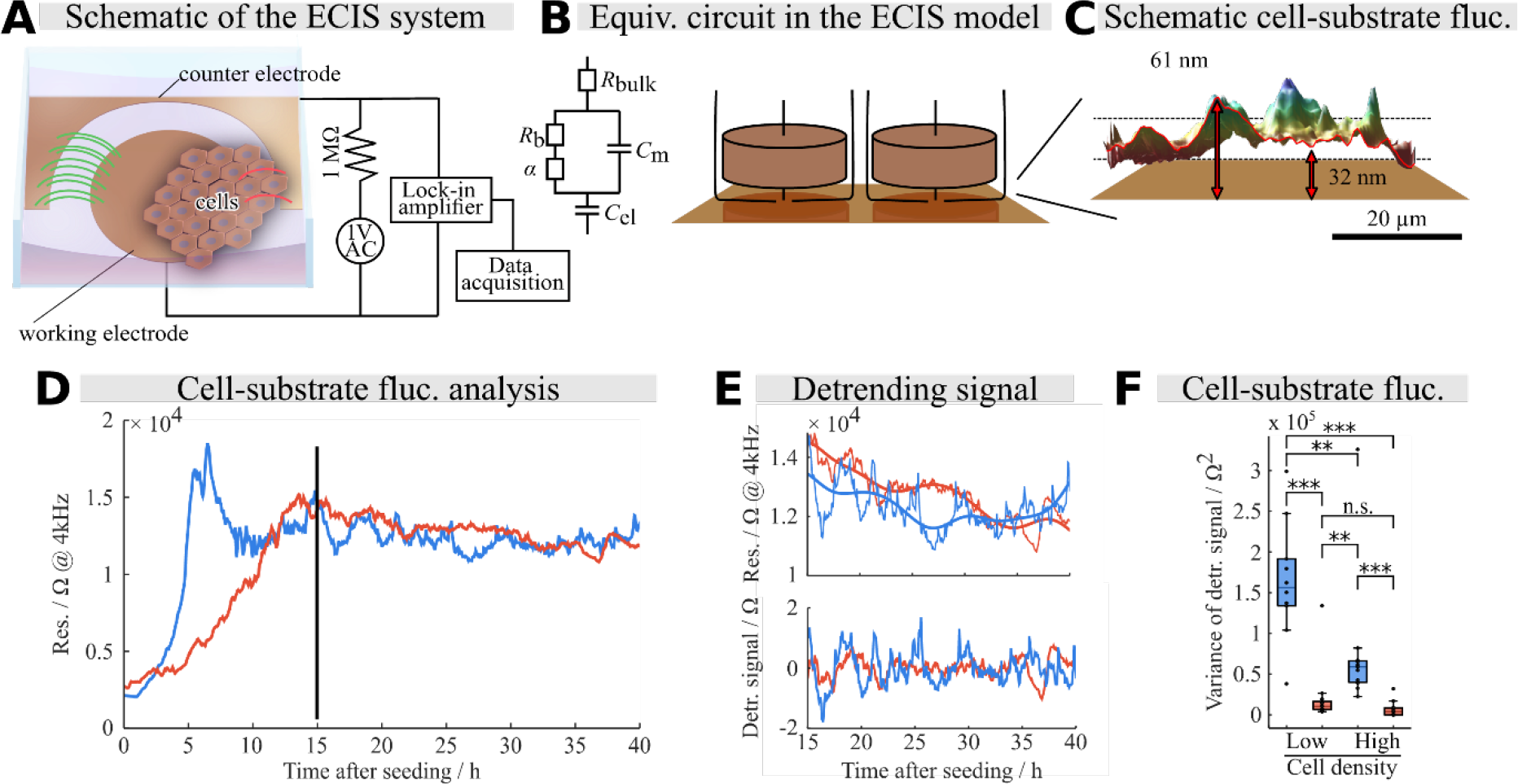
Electrical cell-substrate impedance sensing of cell shape fluctuations. A) Electrochemical setup including a small working electrode hosting the monolayer and a large counter electrode attached to a lock-in amplifier and frequency generator. B) Simplified equivalent circuit describing the impedance response of a confluent monolayer on the electrode together with the envisioned current flow between cells and in between the cleft between electrode and cells. C) MIET image of a MDCKII cell (membrane stained) in contact with a gold surface illustrating the microstructure of the cleft. D) Representative resistance fluctuations of WT and dKD monolayers measured at 4 kHz. After the point of confluency (black line) cell-substrate distance fluctuations of the cells were analyzed. E) Detrending of the signal using empirical mode decomposition (EMD). Top: Trimmed data from D and overall trend (thicker lines). Bottom: Detrended signals after subtraction of the low-frequency overall trend. F) Detrended variance analysis of cell-substrate distance fluctuations measured with ECIS.

Furthermore, we also assessed cell densities of the monolayers at different time points using the cell segmentation software Cellpose.^30^ At elevated cell densities, we expect the mobility of cells to diminish due to jamming as cells adopt a more compact shape, i.e., minimal perimeter at a given area.^31,32^ We found that both cell lines initially have almost identical cell densities at the point of confluence (WT: 1.8×10^5^±2.6×10^4^ cells/cm^2^, dKD: 1.9×10^5^±2.4×10^4^ cells/cm^2^, at 5 h; Figure 1C) after which the cell density increases over time for both cell lines (at 20 h: 3.6×10^5^±5.4×10^4^ cells/cm^2^ (WT), 3.8×10^5^±7.6×10^4^ cells/cm^2^ (dKD)). Furthermore, the probability density of cell areas in WT and dKD cells (SI-Figure 1E+1F) show similar distributions. After 40 h (in a jammed state), however, the cell density of dKD cells is about twofold higher than at the 20-hour interval (8.9×10^5^±2.1×10^5^ cells/cm^2^). In contrast, the cell density of WT cells increases only moderately after jamming (at 40 hours: 5.1×10^5^±1.3×10^5^ cells/cm^2^).

### Cluster sizes correlate with lateral velocities

A cluster size analysis of the cell monolayers was carried out to quantify spatial coherency of the lateral velocity as this manifests the collective nature of migration (Figure 1D+1E). Early (between 5 and 10 h after seeding) and jammed (between 30 and 35 h after seeding) phases of the observation period were examined. Cells moving in the same direction are gathered into a cluster, with larger clusters indicating a more pronounced coherent movement. Generally, WT cells (1296±245 µm^2^) form bigger clusters than dKD cells (921±188 µm^2^) in the early phase (Figure 1I, upper plot). However, the average cluster sizes decrease over time (for 15-20h after seeding (“middle”) see SI-Figure 3A), with WT cells converging to 667±482 µm^2^ and dKD (650±369 µm^2^) assuming similarly small clusters (Figure 1I, lower plot). We further analyzed the shape of clusters finding that the eccentricity of WT and dKD clusters remains constant over time (WT clusters: early: 0.30±0.03; middle: 0.31±0.02; jammed: 0.31±0.03; dKD cluster: early: 0.31±0.02; middle: 0.30±0.02; jammed: 0.31±0.02; SI-Figure 3C), while the aspect ratio (perimeter/area^0.5^) decreases only for WT clusters (early: 5.23±0.41; middle: 5.17±0.54; jammed: 4.88±0.58) not for dKD clusters (early: 4.87±0.24; middle: 4.96±0.27; jammed: 4.88±0.37; SI-Figure 3D). Interestingly, the number of cells per cluster increases in both cell lines in the non-jammed state (SI-Figure 3E) over time, while the cluster sizes in general decrease. This leads to the conclusion that local jamming occurs over time. To quantify this, the velocity maps were re-examined with overlays of the cluster analysis (SI-Figure 3F). In both cell lines, larger clusters displayed higher local velocities, while smaller clusters with more jammed cells showed lower local velocities. Correspondingly, the decay of the lateral velocity correlation (Figure 1F, upper plot) in early phases was less for WT monolayers compared to dKD. The correlation length found for WT cells was 62.6±17.5 µm, significantly higher than for dKD cells (45.6±4.8 µm; Figure 1G, upper plot). Similar to the average velocity, the correlation length dropped in the jammed state to the same level for both cell lines (Figure 1G, lower plot).

**Figure 3:**
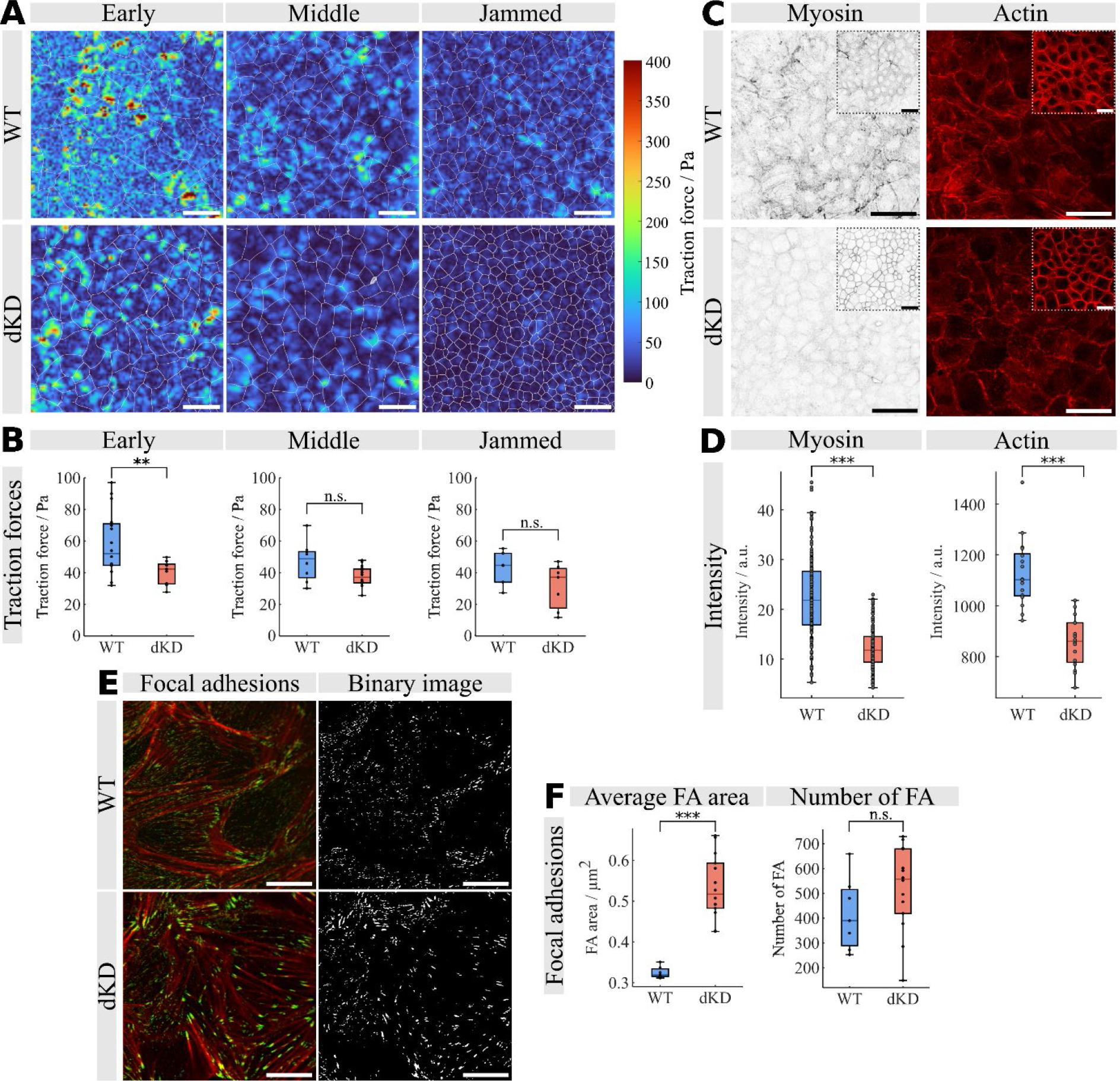
Traction force microscopy, actomyosin architecture, and quantification of focal contacts. A) Representative traction force maps of WT and dKD monolayers with cell outlines (white) obtained from Cellpose. Scale bars: 40µm. B) Average traction forces of monolayers at three different time points shown in A). C) Representative fluorescence images of WT and dKD monolayers seeded on PAA gels (basal side, inverse contrast) showing actin (Phalloidin) and myosin-II. Upper corners show corresponding images of the apical side. Scale bar: 40 µm. D) Fluorescence intensities of stained myosin and actin, respectively. E) Representative fluorescence images of WT and dKD monolayers (basal side) seeded on glass showing actin (Phalloidin, red), paxillin (green), and the generated binary images. F) Quantification of focal adhesions (FA).

Additionally, we analyzed mean square displacements (MSD) for monolayers during early phases (Figure 1F, lower plot) to assess their rearrangement capabilities. Generally, WT cells exhibit higher MSDs than dKD cells (Figure 1H). The MSD (MSD ∝ *τ*^α^) allows us to deduce cell rearrangement activity from the scaling exponent (α) for different cell lines and conditions. An α value of 2 suggests linear movement, while α = 1 indicates Brownian motion, and smaller values imply enhanced interaction with neighboring cells leading to subdiffusive motion. The MSD exponent is considerably higher for WT (1.23±0.04), indicative of superdiffusive motion compared to dKD (1.02±0.04). As cell density increases, the exponent decreases for both samples (Figure 1H, lower plot; SI-Figure 2B), suggesting decreased mobility as cells approach the jamming limit, aligning with previous research.^10,22^ However, the exponent in WT cells remains always significantly higher than for dKD cells (WT: 1.06±0.12; dKD: 0.81±0.10) even in the jammed state where the PIV method fails to generate useful data. We attribute this inherent difference in mobility to the contractile nature of dKD cells paired with an altered interaction with the substrate as discussed below.

### Changes in cell-substrate distance fluctuations mirror lateral average velocity changes

Cellular motion extends beyond the collective lateral 2D migration of cells; it also comprises cell membrane fluctuations with respect to the adhesion surface (cell-substrate distance fluctuation), essential for the dynamic opening and closing of adhesion bonds formed with the ECM. An unobtrusive technique that can probe the space between cells and their underlying surface employs electrical cell-substrate impedance sensing (ECIS). This method utilizes ultrasmall electrodes to guarantee that the measured resistance is determined predominately by the adherent cells. The impedance spectra typically cover a frequency range of 1 Hz to 100 kHz and can be described by a combination of resistors and capacitances representing the dielectric properties of the cells and interfaces to the substrate or adjacent cells, respectively. The typical impedance spectra of WT and dKD monolayers (SI-Figure 4B) can be divided into five distinct regimes. At the lowest frequencies, electrode charging predominantly affects the complex impedance (1). After this dispersion an intermediate regime follows at frequencies approximately between 100 Hz - 4 kHz, where the cell-substrate interface becomes significant due to restricted current flow between the cells and the substrate (2). Next, a purely ohmic behavior is observed, which mainly represents the transepithelial resistance between the cells (3). This is followed by a second strong dispersion resulting from the dielectric properties of the cell membranes (4). At high frequencies > 100 kHz, the resistance of the medium dominates the response (5).^33–35^ As we were primarily interested in cell-substrate fluctuations as a function of time, we chose to record impedance fluctuation at 4 kHz and below (Figure 2A-C, SI-Figure 4C). At these frequencies, the normalized impedance of both cell lines is almost identical (SI-Figure 4C), with the difference in fluctuation amplitude between the cell lines increasing at lower frequencies (see 1 kHz). Confirmed by optical inspection, confluence of the cell layer was determined based on the point at which the overall impedance reached a plateau (black line in Figure 2D). The following 25 h were subsequently analyzed (Figure 2E, upper plot). These trimmed data sets contain both cell-substrate distance fluctuations and an overall long-term trend due to the maturation of the monolayer, which was removed by empirical mode decomposition (EMD, SI-Figure 5), as we are mainly interested in short-term fluctuations of the cleft size between the cell and the electrode.^36–38^ Subtracting the lowest-frequency modes from the original signal results in a detrended signal (Figure 2E, lower plot), which can be used to obtain the variance and temporal persistence (Figure 2F, SI-Figure 6C).^39^ While the variance corresponds to the fluctuation amplitude, the scaling exponent of the detrended fluctuation analysis (DFA), α, reflects the time series’ self-correlations. At frequencies below 4 kHz, the WT cell layers show a higher variance (3.2×10^5^±1.0×10^5^ Ω^2^) of the cell-substrate distance fluctuations compared to dKD cells (8.0×10^4^±4.0×10^4^ Ω^2^) and a higher α-value (WT: 1.47±0.30, dKD: 1.16±0.13). This indicates that more substantial, actively driven cell-substrate distance fluctuations occur in WT cells than in dKD cells. At the same time, the larger α-values reflect the emergence of temporal persistence correlations (persistent motion) more pronounced for WT cells. α-values reflect the memory (autocorrelation) of the time series as detailed in the SI. We also compared the cell densities in the ECIS chambers with the cell densities used for the lateral velocity measurements. Both WT (2.6×10^5^±2.0×10^4^ cells/cm^2^) and dKD (2.5×10^5^±1.2×10^4^ cells/cm^2^) have similar cell densities at the beginning of the ECIS measurement as well as in the lateral velocity measurements after 10 h (WT: 2.3×10^5^±2.5×10^4^ cells/cm^2^, dKD: 2.4×10^5^±3.0×10^4^ cells/cm^2^). Additionally, we performed impedance measurements with a doubled cell density to examine the cell-substrate distance fluctuations of denser monolayers (Figure 2F). Both cell lines display smaller impedance variances compared to the less dense monolayers, going hand in hand with the findings of decreasing average lateral velocity with increasing cell densities. We found that the average lateral velocity and cell-substrate distance fluctuations are highly correlated for WT cells: Changing the extracellular matrix composition to vary the substrate adhesions concomitantly alters cell-substrate distance fluctuations and lateral velocity. Conversely, dKD cells show almost no influence of substrate coating on cell-substrate distance fluctuations, resulting in no correlation to the average velocity. The data (SI-Figure 4A) also confirms that cell-substrate distance fluctuations dominate over resistance fluctuations originating from variations of the cell-cell contacts.

**Figure 4.**
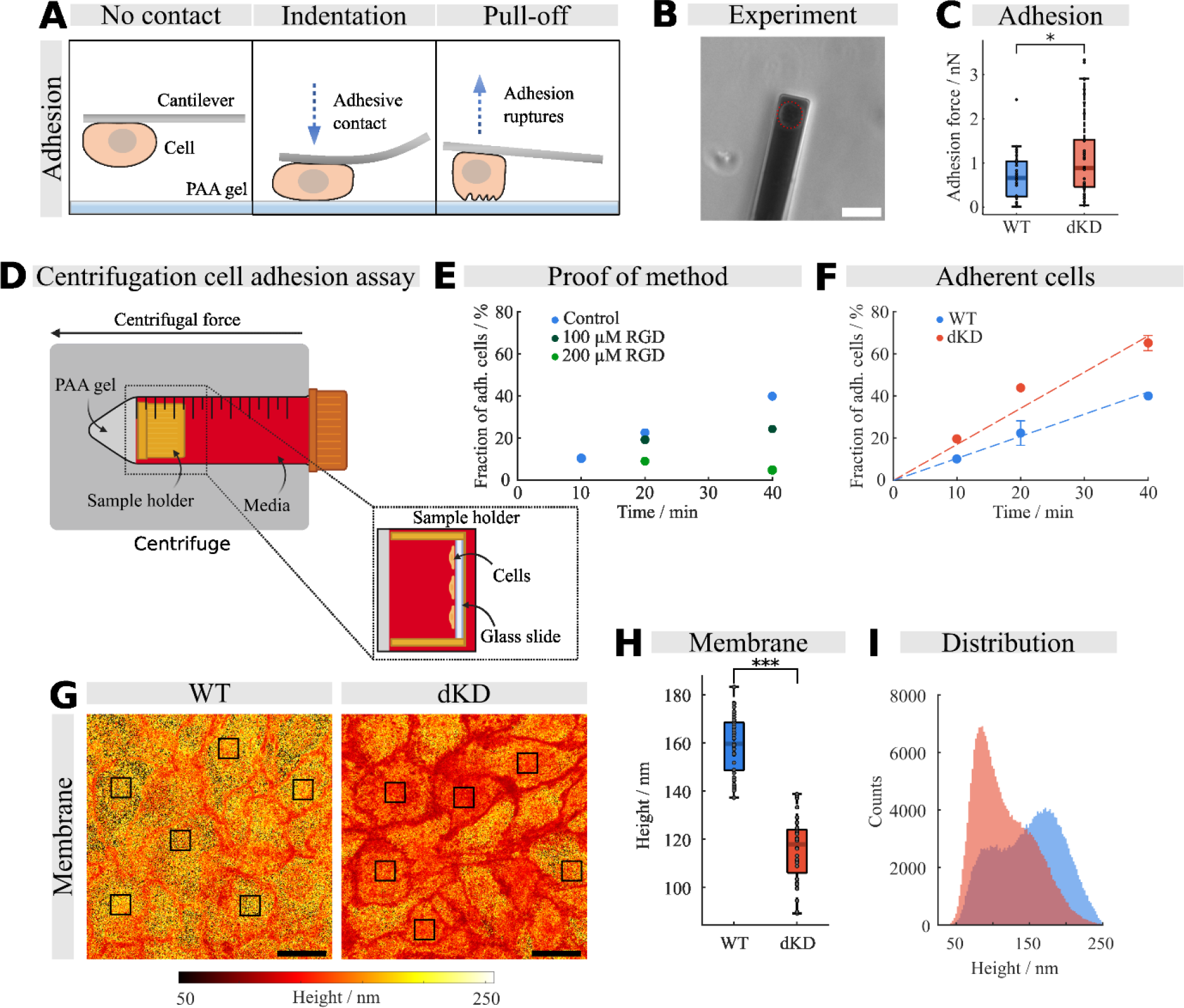
Cell-substrate adhesions correlate with average membrane-substrate distances. A) AFM measurement of cell-substrate adhesion forces. The cell attached to the cantilever is brought in contact with a polyacrylamide (PAA) gel for 150 s. Afterward, adhesion is measured by disengaging the cell from the gel. B) Optical micrograph of a cell attached to the tipless cantilever. Scale bar: 50 µm. C) Average cell-substrate adhesions (max. adhesion force) on a 1.6 kPa stiff PAA gel. D) Scheme illustrating the centrifugation assay. E) Proof-of-principle measurements for the centrifugation assay. Different concentrations of the peptide RGD were used to inhibit cell-substrate adhesion. F) Fraction of cells still attached to the surface after centrifugation as a function of time after cell seeding. G) Representative height (distance between the gold substrate and basal membrane) maps obtained from MIET microscopy. For further analysis, areas in cell centers were selected (black rectangles), and the corresponding height values are shown in H). Scale bar: 20 µm. I) Representative height distributions of a confluent cell layer (100 µm x 100 µm).

**Figure 5.**
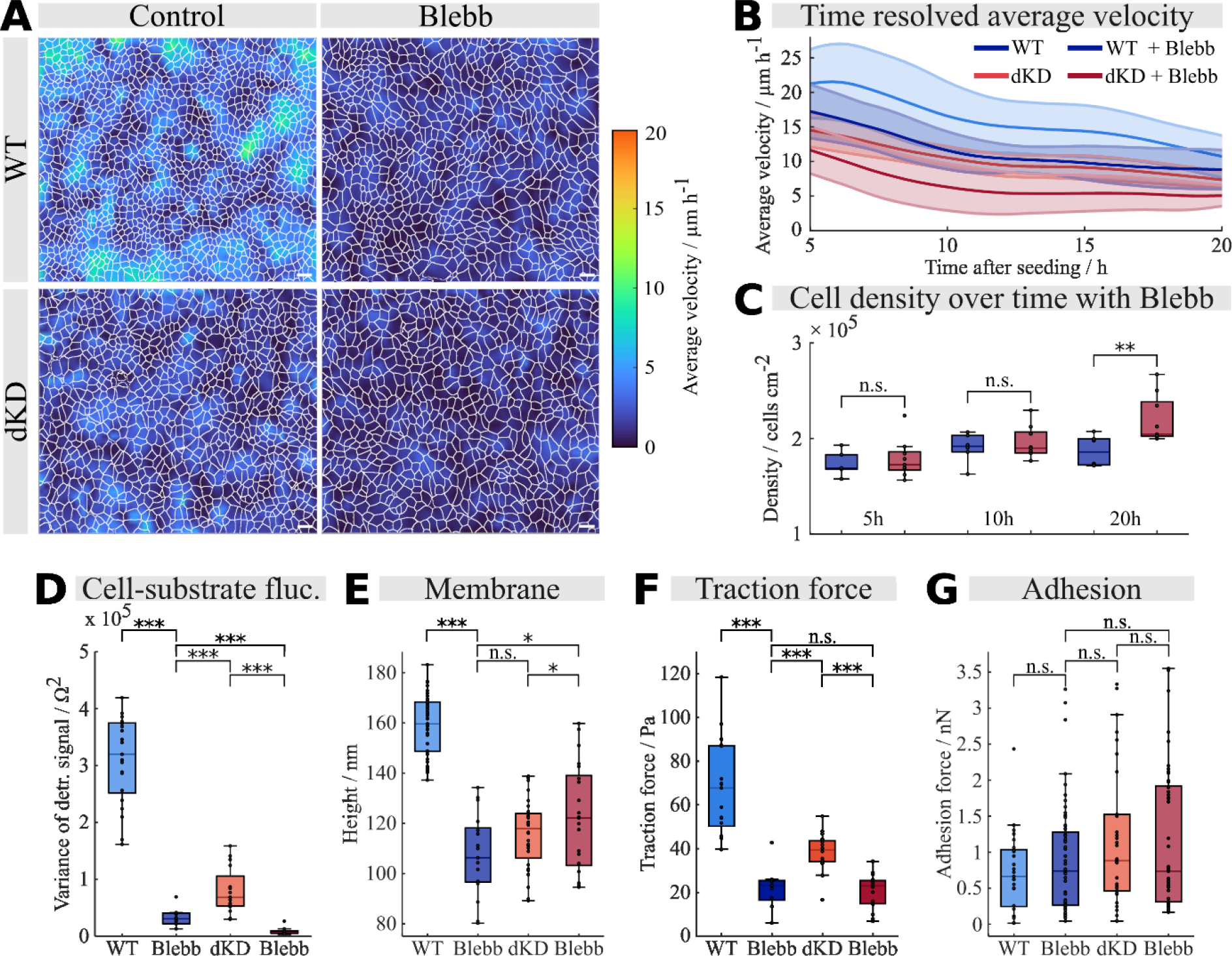
Impact of myosin-II on cellular motility. A) Representative velocity maps generated from the corresponding velocity vectors obtained from PIV. Cell outlines (white) were obtained from Cellpose. Scale bars: 40 µm. B) Average velocities of WT and dKD monolayers treated with 50 µM Blebb compared with untreated monolayers. C) Cell densities of Blebb-treated cells obtained from Cellpose. D) Detrended variance analysis of cell-substrate distance fluctuations measured with ECIS. E) Average height values of the basal membrane from the gold surface calculated from selected cell centers of MIET microscopy images. F) Average traction forces of confluent monolayers 10 h - 20 h after seeding. G) Average cell-substrate adhesion forces (max. value) measured by AFM.

**Figure 6.**
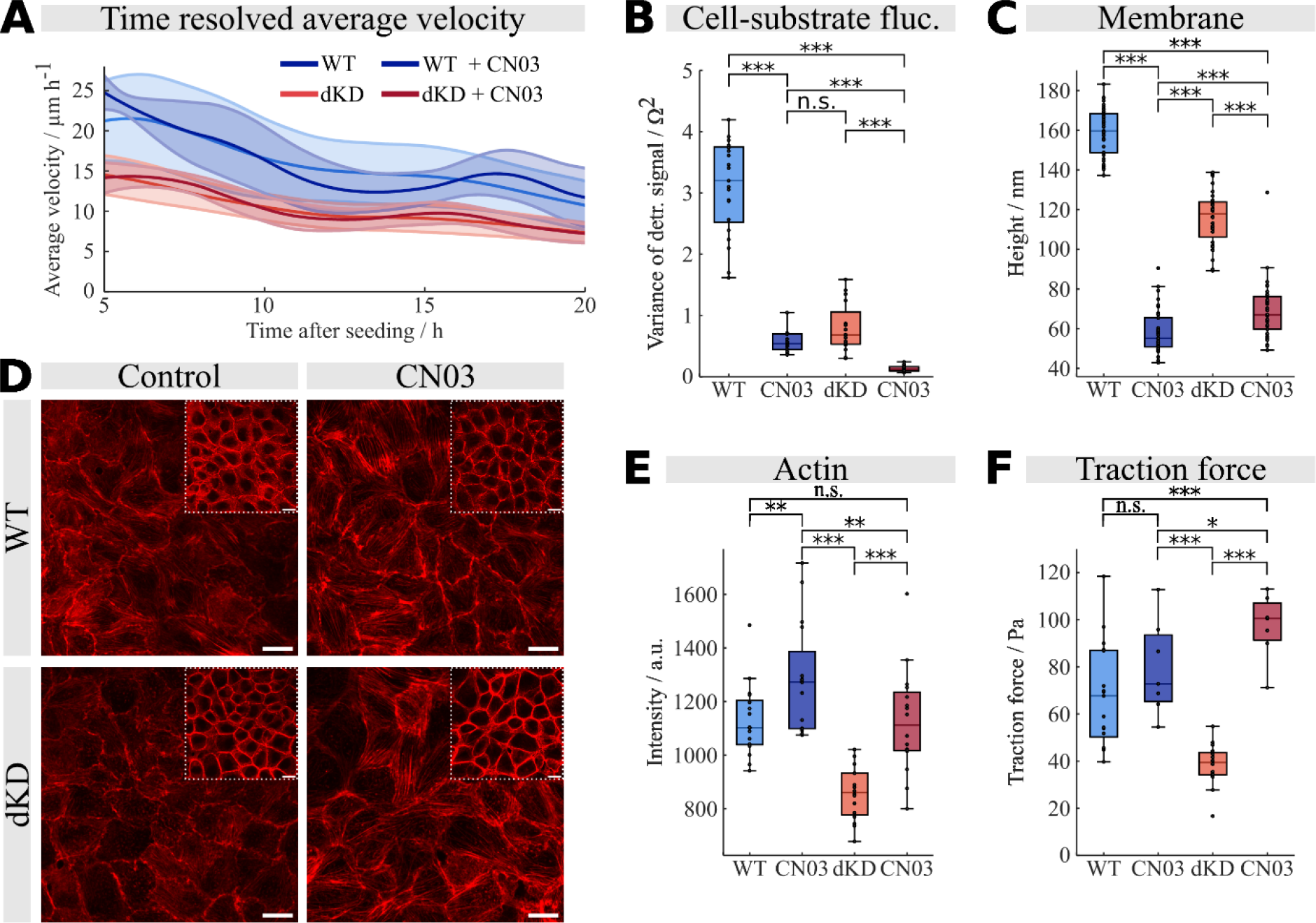
Impact of Rho activation on cellular motility. A) Average velocities of WT and dKD monolayers treated with 1 µg/mL of Rho activator CN03 compared with untreated monolayers. B) Detrended variance analysis of cell-substrate distance fluctuations measured with ECIS. C) Average height values of the basal membrane from the gold surface calculated from selected cell centers of MIET microscopy images. D) Representative fluorescence images of WT and dKD monolayers seeded on PAA gels (basal side) showing actin (Phalloidin). Upper corners show the corresponding images of the apical side. Scale bars: 15 µm. E) Fluorescence intensities of actin. F) Average traction forces of confluent monolayer 10 h - 20 h after seeding.

In summary, dKD cells exhibit substantially lower average lateral velocity and diminished amplitude and persistence of cell-substrate distance fluctuations than WT cells. The reduction in three-dimensional mobility persists within the 25-hour period following cell confluency, leading up to the cells’ jamming (representing a glassy state with substantially reduced mobility). The reduced lateral motion of dKD cells is attributable to the contractile nature of the apical side. Hence, dKD cells can enter an exceedingly dense configuration in the jamming limit. Conversely, reasons behind the variations in the distance fluctuations between the cell and the substrate are yet to be elucidated. To understand this, we need to measure the involved forces, including adhesion and traction forces, the precise cell-substrate distance, and quantify the participating cytoskeletal proteins.

### Average traction forces decrease with decreasing average velocity

We hypothesize that variations in the distance between cells and substrate are a result of the cell’s inherent actomyosin contractility. Such fluctuations are likely to be reflected in cellular traction forces (Figure 3A) and immunostaining of myosin (vide infra). We found that WT monolayers generated higher average traction forces (58±20 Pa) compared to dKD monolayers (40±7 Pa) 10 hours (cell density: 1.7 - 2.6×10^5^ cells/cm^2^) after reaching confluence (early, Figure 3B). Average traction forces decrease with decreasing lateral velocity of the monolayers. In the middle phase, WT monolayers exert average traction forces of 47±13 Pa, while dKD show average traction forces of 38±6 Pa (cell density: 2.8 – 4.5×10^5^ cells/cm^2^). The average traction forces drop further in the jammed phase (cell density: 5.5 – 8.0×10^5^ cells/cm^2^), with a more significant decrease in traction forces in the WT cell line (42±10 Pa) than for dKD cells (32±13 Pa) also concurrent with the drop in lateral velocity.

We also performed traction force experiments in the presence of Mitomycin C (MitoC) to separate the impact of longer incubation time and density increase by arresting cells before entering the S-Phase of the cell cycle and effectively blocking proliferation.^40,41^ Low concentrations of MitoC (10 μg/mL) have been shown to cause proliferation inhibition without changing growth or morphology while causing a slight increase in apoptosis as a side effect.^42^ Monolayers were treated with MitoC after 20 hours and incubated for an additional 24 hours. The average traction forces of these MitoC-treated monolayers are slightly higher than untreated monolayers 20 hours after the point of confluence (SI-Figure 7). This proves that the cell layers do not reduce the average traction forces due to longer incubation time, but rather when entering the jammed state as the cell density increases.

### dKD cells show contractility switch from basal to apical

Previous studies have shown that cells lacking ZO-1/2 are more contractile by expressing a pronounced apical actomyosin ring, while WT cells do not show any upregulation.^21,22^ It therefore needs further investigations to understand why traction forces are reduced in contractile cells and why cell-substrate distance fluctuations are diminished. Consequently, we used confocal fluorescence microscopy to visualize both F-actin and myosin light chain kinase, essential proteins of the basal actomyosin skeleton, involved in generating cellular traction forces.^43,44^

The increased apical actomyosin ring can also be seen in dKD cells cultured on PAA substrates (small figures in the right upper corners in Figure 3C). However, we observed structural changes in the actomyosin skeleton on the basal side of the monolayers. dKD cells revealed, on average, 46% lower myosin intensity and 24% lower actin intensity than WT cells (Figure 3D) indicating decreased activity of myosin and actin on the basal side, correlating with the differences we found in traction force maps. As a result, an increase in myosin and actin activity on the basal cell side is associated with higher average traction forces and higher fluctuation amplitudes of the cell-substrate distance in monolayers of WT cells.

In the next section, we address the cells’ adhesion to the substrate, comprising the dynamic adhesion strength and the maturation of focal contacts.

### ZO-1/2 dKD cells show increased cell-substrate adhesion

First, we employed confocal microscopy to visualize paxillin expression (Figure 3E) to assess the number and size of focal adhesions. Our findings revealed that the average size of focal adhesions in dKD monolayers (0.53±0.08 µm^2^) was higher than in WT monolayers (0.32±0.01 µm^2^), while the number of focal adhesions (WT: 417±147, dKD: 522±170) was largely preserved (Figure 3F). The findings indicate that the diminished motility of confluent dKD cells leads to the formation of larger focal adhesions or vice versa. Regardless of cause and effect, larger focal adhesions display decreased dynamics. This observation aligns with previous findings in individual cells, where it was noted that cell movement decelerates when the size of focal adhesions surpasses a certain threshold.^13^ Therefore, we propose that this size (and, therefore, potential dynamics) difference also affects cell-substrate adhesions and, eventually, cell-substrate distance fluctuations.

Second, we used single-cell force spectroscopy (via AFM) to examine the dynamic bonding between the substrate and the cells (Figure 4A). A solitary cell was affixed to a tipless cantilever and positioned in contact with a PAA gel for a specified duration and load force to facilitate the formation of cell-substrate adhesions (Figure 4B). Subsequently, the cell was disengaged from the surface, and the maximum force required for detachment was documented. We found that WT cells exhibited slightly lower cell-substrate adhesion forces after a contact time of 150 s compared to dKD cells (Figure 4C). However, as this method is extremely precise but cannot be used for longer incubation times due to the elevated forces once adhesions start to mature, we also used a centrifuge assay to quantify the fraction of cells that withstand rupture from the surface during centrifugation (Figure 4D).^45,46^ This method applies detachment forces to numerous cells. With increasing cell numbers remaining on the substrate after experiencing centrifugal forces, stronger cell-substrate adhesion forces per area are assumed. To validate the method, the cells were incubated with a short peptide RGD (arginine-glycine-aspartic acid) at elevated concentrations (Figure 4E). It is known that certain integrins are blocked by RGD peptide, a known binding sequence of extracellular matrix proteins such as fibronectin or collagen.^47,48^ With increasing RGD concentration, the fraction of adherent cells decreases, demonstrating the specificity of the centrifugation assay for integrin-mediated cell-substrate adhesions.

We observed higher fractions of adherent cells in dKD samples (19.5±4.2 %) compared to WT cells (10.4±2.2 %) after 10 minutes of incubation followed by centrifugation, suggesting that dKD cells produce larger adhesion forces per area. The fraction of cells remaining on the substrate increases with longer incubation times of 20 and 40 minutes for both cell lines. AFM and centrifugation assays both demonstrate that dKD are more firmly attached to the substrate than WT cells, which, in contrast, show larger traction forces. As a result, the formation of ligand-receptor bonds within the contact area leads to increased stick-slip frictional forces in dKD cells.^49^ Taken together, we assume that elevated myosin activity is responsible for the reduced number of closed bonds in WT cells. This is further investigated by measuring the distance between the cells and the substrate and abolishing myosin activity (*vide infra*).

### Cell-substrate distance measured by metal-induced energy transfer (MIET) microscopy

Less traction force and higher adhesion force should result in an altered cell-substrate distance. To quantify this, we performed membrane height studies with both cell lines using MIET microscopy, providing unprecedented nanometer z-resolution, allowing a spatial inspection of the cleft between the basal cell side and the gold substrate.^50–54^ For membrane height quantification, we selected six rectangles in the center of a cell from each measurement (Figure 4G). The extracted points obtained from each of these rectangles were averaged. We found that the WT cells show an average height of (159±12 nm), while dKD cells were closer to the substrate, with an average height of (116±14 nm; Figure 4H+4I). This finding matches the stronger adhesion forces and reduced cell-substrate distance fluctuations (*vide supra*).

### Actomyosin activity governs the dynamics of individual cells and thereby steers collective migration

Based on our experiments, we deduce that the observed decrease in basal contractility and the fluctuation of cell-substrate distance in confluent dKD cells, relative to WT cells, is attributable to diminished myosin activity. Hence, we conducted experiments in the presence of Blebbistatin (Blebb), a selective inhibitor of myosin-II, to stall the overall contractility.^55^ We found a decreased collective lateral velocity of confluent monolayers (both cell lines) treated with 50 µM Blebb over time (Figure 5A+5B). The average velocity of WT cells decreased from 17±3.8 µm/h after 5 hours to merely 9.2±3.1 µm/h after 20 hours. In contrast, dKD cells were slower throughout the entire observation time. Their velocity was 12±3.4 µm/h after 5 hours and decreased below the detection threshold of 4.8±0.7 µm/h after 20 hours. The cell density remains almost constant during the observed period (Figure 5C).

Concomitantly, cell-substrate fluctuations drop substantially in response to myosin-II inhibition (Figure 5D). The WT monolayers still display a slightly higher variance of the cell-substrate-fluctuations (3.3×10^4^±1.6×10^4^ Ω^2^) compared to dKD cells (9.1×10^3^±6.6×10^3^ Ω^2^) in the presence of Blebb, which are probably just thermally excited membrane/cortex undulations as their correlation (α-value) also drops. We conclude that reducing the basal contractility of the actomyosin cytoskeleton impairs the cells’ ability to perform essential vertical movements, which are necessary for generating traction forces, ultimately inhibiting collective cell migration. The remodeling of the actin cortex to reinforce the apical ring at the expense of basal fibers is held responsible for that.

Interestingly, as the cells’ contractile ability is impaired, both WT and dKD cells reveal a similar lateral velocity. This suggests that the initial variations in cell motility and dynamics arise from the contractile machinery of the basal actomyosin cytoskeleton. Suppressing myosin-II activity in WT cells eliminates these differences, leading to a convergence in behavior between WT and dKD cells. The importance of basal contractility is further supported by measuring the distance between the basal membrane and the substrate using MIET microscopy (Figure 5E). Following Blebb treatment, the average height of the basal membrane in WT cells is 116±23 nm. Consequently, this aligns with the basal membrane height of dKD cells without Blebb treatment (116±14 nm). Inhibition of myosin-II in dKD cells does not affect the distance between the substrate and membrane, remaining very similar at 126±21 nm. This indicates that the larger distance between the substrate and basal membrane in WT cells compared to dKD cells is actively driven by the basal actomyosin cytoskeleton fostering also the collective migration.

A similar trend is also found for the traction forces exerted by the cells on the substrate. Both WT (22±10 Pa) and dKD (20±8.0 Pa) cells show similar traction forces after treatment with Blebb (Figure 5F), substantially reduced compared to untreated control cells (traction forces between 10 h and 20 h were averaged). This observation emphasizes that reduction of the basal contractility of WT cells causes a comparable phenotype to untreated dKD cells in which basal contractility is reduced, leading eventually to reduced cell-substrate fluctuations.

Interestingly, cell-substrate adhesion forces are only slightly influenced by the reduction of contractility (Figure 5G). We noted a marginal increase in adhesion forces in WT (0.9±0.7 nN) and dKD (1.6±1.0 nN), respectively, showing again that adhesion forces are not correlated with traction forces.

We deduce that the effective interaction between the substrate and the basal cell membrane is diminished in WT cells due to increased contractility of the basal actomyosin network compared to dKD cells. This results in elevated cell-substrate fluctuation amplitude, temporal persistence, increased overall traction forces and decreased cell-substrate adhesion enabling cells to move more rapidly and coherently. Following treatment with Blebbistatin, WT cells and dKD cells exhibit similar characteristics, with both types of cells demonstrating a decrease in three-dimensional fluidity and traction forces.

### Rho activation reinforces the basal actomyosin cytoskeleton

We used the Rho activator CN03 (1 µg/mL), which is known to increase the number and order of stress fibers,^10^ to rescue the reduced basal contractility of dKD cells. While the average lateral velocity does not change over time compared to untreated monolayers (Figure 6A), the cell-substrate distance fluctuations are dramatically reduced (Figure 6B). The variance of CN03-treated WT monolayers (5.9×10^4^±2.2×10^4^ Ω^2^) dropped to the level of untreated dKD cells (8.0×10^4^±4.0×10^4^ Ω^2^, Figure 1F), while cell-substrate fluctuations decrease even further in CN03-treated dKD monolayers (1.3×10^4^±5.1×10^3^ Ω^2^).

Measurements with MIET microscopy show that the basal membranes of both cell lines treated with CN03 are, on average, substantially closer to the substrate (Figure 6C). WT basal membranes (59±11 nm) are more than twice as close to the substrate as untreated WT cells (159±12 nm), and dKD cells also show a reduced distance from the substrate (treated: 70±14 nm, untreated 116±14 nm).

This is supported and rationalized by fluorescence images of actin on the basal side before and after treatment with CN03 (Figure 6D). In response to Rho activation, the basal intensity of actin in dKD monolayers (1124±196 a. u.) reaches the same level as the untreated WT monolayers (1129±132 a. u.), while the CN03-treated WT monolayers (1284±203 a. u.) show only a small increase in actin intensity (Figure 6E). This means we can reestablish the loss of basal contractility in dKD monolayers by administering CN03.

Subsequently, we performed traction force experiments with CN03-treated monolayers. We averaged traction force data sets collected between 10 h and 20 h after seeding (Figure 6F). We found that, on average, CN03-treated dKD monolayers generate higher traction forces (97±14 Pa) than untreated dKD cells. The traction forces of WT monolayers are also (albeit insignificantly) affected (CN03 treatment: 79±20 Pa). The data from both sets reveal a consistent relationship where increased actin intensity is associated with stronger traction forces.

These findings indicate that Rho activation leads to more stress fibers, increasing basal contractility, leading to higher traction forces, essentially ironing out cell-substrate distance fluctuations and pull the cells even closer to the substrate. However, to our surprise, the cells maintain their collective lateral velocity, with neither the WT cells slowing down nor the dKD cells gaining speed. This implies that the cells compensate for decreased vertical dynamics by stronger traction forces that overcome the increased interaction with the substrate.

## Discussion

The concept of collective migration surpasses the mere coordinated movement of individual cells.^2^ Thus, the crux of collective cell migration lies in cells’ capacity to reposition themselves relative to their neighbors with minimal substrate interaction. The shape of cells expedites cellular reorganizations, as those with elongated forms and larger perimeters can more readily glide past neighboring cells within the cell layer.^1,10,32,56–59^ While it is believed that cortical tension and adhesion at the periphery of each cell predominantly regulate cell perimeter, experimental investigations into this hypothesis have yielded contradictory outcomes.^10^ The conflicting results have recently been resolved by examining forces, cell perimeters, and motion of an epithelial monolayer as a function of cell density.^10^ The authors observed a simultaneous decrease in all three factors with elevated cell density or the inhibition of cell contraction. Interestingly, the augmentation of tractions was identified as a factor capable of counteracting the influence of density on cell shape and rearrangements. Consequently, their study shifts the emphasis away from cell-cell contacts, highlighting cell-substrate traction as a primary physical determinant governing shape and motion in collective cell migration.

Here, we extend this picture by adding cell-substrate distance fluctuations using two cell lines that differ substantially in collective migration due to their intrinsic contractility: MDCKII wild-type (WT) and MDCKII ZO-1/ZO-2 double knock-down (dKD) cells. Loss of these two ZO proteins in tight junctions (TJs) leads to an imbalance in the actomyosin skeleton on soft polyacrylamide gels. A thick, highly contractile actomyosin ring is formed on the apical side in dKD cells, sacrificing basal contractility, which is especially pronounced in later stages when jamming occurs. It is assumed that TJs exert negative mechanical feedback on the actomyosin cytoskeleton of individual cells in a cell layer.^16^ This feedback function helps to prevent excessive contraction and elongation of the cells and, therefore, maintains tissue integrity. In several TJ-KO cell lines, it has been observed that this feedback loop is missing, leading to uncontrolled apical contraction.^16,22^ In contrast, little is known about the basal actomyosin skeleton in TJ-KO cells. We found pronounced actin and myosin gradients in dKD cells, suggesting reduced basal contractility, as recently reported for glassy substrates.^26^ The altered basal actomyosin skeleton in dKD cells explains the reduced average traction forces in dKD monolayers. Monolayers with higher density exerted lower traction forces on average regardless of the cell type, as also found by Saraswathibhatla et al.^10^ Here, we could rule out that increased proliferation is responsible for the observed alteration of traction forces.

Furthermore, we show that confluent dKD monolayers exhibit reduced collective migration velocity paired with lower traction forces, larger cell-substrate adhesion, and a more compact shape. We consistently confirm average velocity decreases in both cell lines as cell density rises.^31,32,57^ Saraswathibhatla et al. identified analogous correlations when examining cell layers with varying densities. Monolayers with higher cell density exhibited lower traction forces compared to low cell density.^10^

In the PIV data, we also observed a limiting velocity of 4 µm/h set by the method and consistent with earlier findings.^31^ Even beyond this limit, cell density in dKD cell layers continues to increase, while density is preserved in WT cells. This effect can be explained by the role of ZO proteins in controlling proliferation through cell cycle arrest.^60,61^ Hence, when TJs are depleted, cells lose contact inhibition. Moreover, dKD cells show smaller cell clusters with coordinated movement, a lower scaling exponent of the MSD, and shorter correlation lengths indicative of reduced cell-cell force transmission. Importantly, WT cells display substantially larger cell-substrate distance fluctuations allowing the cell to reduce interaction with the substrate and respond swiftly to mechanosignalling. Fluctuations are found to be responsible for increasing the average cell-substrate distance and, therefore, allowing the cells to remain in an out-of-equilibrium state that enables them to move more collectively at elevated speed.^62^ Sabass and Schwarz showed how cell-substrate friction depends on the number of bonds, their dynamics, and intracellular relaxation.^63^ In this state, only slight variations in control parameters, such as on/off rates of adhesion molecules, lead to significant changes in macroscopic quantities characterizing the system. This permits the cell to bridge length scales on short time scales by small variations in contractility. Elevated cell-substrate distance fluctuations paired with increased average cell-substrate distance are also found at the onset of the epithelial-to-mesenchymal transition (EMT), where cells enter a state of higher mobility, larger distance from the surface, and more fusiform shape – all parameters allowing the cells to move more rapidly and independently.^52,64,65^ An increase in cell density further reduces basal movement, which aligns with the observed reduced lateral velocity.^52^ It is well known that migration and, therefore also, basal movement of the membrane is significantly influenced by myosin-II.^66^

The short-range force transmission in cell clusters, resulting from cell-matrix adhesions, contrasts with the longer-range force transmission required for processes like collective cell migration. Active regulation of force propagation through spatial coordination of actomyosin contractility is proposed to achieve this longer-range force transmission, as seen in tissue morphogenesis.^67,68^ The reduction in spatial and temporal persistence in confluent dKD cells before jamming results in compromised long-range force transmission, leading to slower and less coherent migration mirrored in smaller clusters with the same velocity. This is further fostered by increased cell adhesion, which inhibits long-range transmission. Conversely, more loosely attached cells can effectively relay forces, as we see with WT cells.^68^

Applying CN03 results in an elevated quantity and reinforcement of stress fibers and consequently augments traction forces, which could maintain lateral velocity even at reduced cell-substrate distance fluctuations. Conversely, using blebbistatin decreases cell-substrate distance fluctuations down to pure thermal noise with no memory, reducing the average lateral velocity of the cell layer. These two experiments support the idea that a counterbalance of traction forces and cell-substrate distance fluctuations enables fast and coherent collective migration of confluent cell layers.

The idea that shape fluctuations might foster 2D velocity has recently been proposed by Manning and coworkers using tissue models.^69^ According to Yanagida et al., sorting of the primitive endoderm from the epiblast in the inner cell mass in early mammalian embryos is attributed to differential cell surface fluctuations rather than static physical parameters.^70^

In summary, our data reveal that cells utilize chemical energy to produce significant cell-substrate distance fluctuations, surpassing thermal levels, which are crucial for achieving and sustaining high lateral cell velocities. These fluctuations elevate the basal cell membrane, and in conjunction with adequate traction forces, facilitate positional changes within the layer. Our research underscores the critical role of stress fibers and myosin in exerting the essential forces at the cell-substrate interface. These forces lead to persistent, active shape fluctuations, propelling motile cells in a manner like active Brownian particles at an increased effective temperature. Ultimately, this dynamic process enables coordinated cell movements, which are disrupted by imbalanced contractility, as observed in ZO1-depleted MDCKII cells or when myosin function is impaired by drugs like blebbistatin.

### Limitations of the study

Investigating the collective migration of cells necessitates exploring an extensive range of parameters, which carries the risk of unintended consequences when disrupting certain functions like myosin activity or applying drugs that compromise the structure of the cytoskeleton. Such side effects may involve the cytoskeleton’s reorganization to offset functional deficits. Additionally, it should be noted that while the data presented here come from a well-established model, they may not universally apply to all types of epithelial cells.

## Acknowledgments

Funding by the DFG (grant JA963/19-1) is gratefully acknowledged. The authors thank Burkhard Geil for helpful discussions on image analysis, segmentation, tracking, and empirical mode decomposition. The authors would also like to acknowledge Jörg Enderlein for his assistance and expertise in MIET, and Jonathan Bodenschatz for his help with the centrifugation assay analysis. Figure 4D was created with BioRender.com.

## Author Contributions

Conceptualization, M.J., A.J., and T.A.O.; Software, M.J., A.T., and P.N.; Investigation, M.J., B.D.W., L.E., A.T., A.R., and A.C.; Resources, A.H., and A.C.; Writing - Original Draft, M.J., T.A.O., and A.J.; Writing - Review&Editing, M.J., T.A.O., and A.J.; Funding Acquisition, A.J.

## Star Methods

### Key resources table

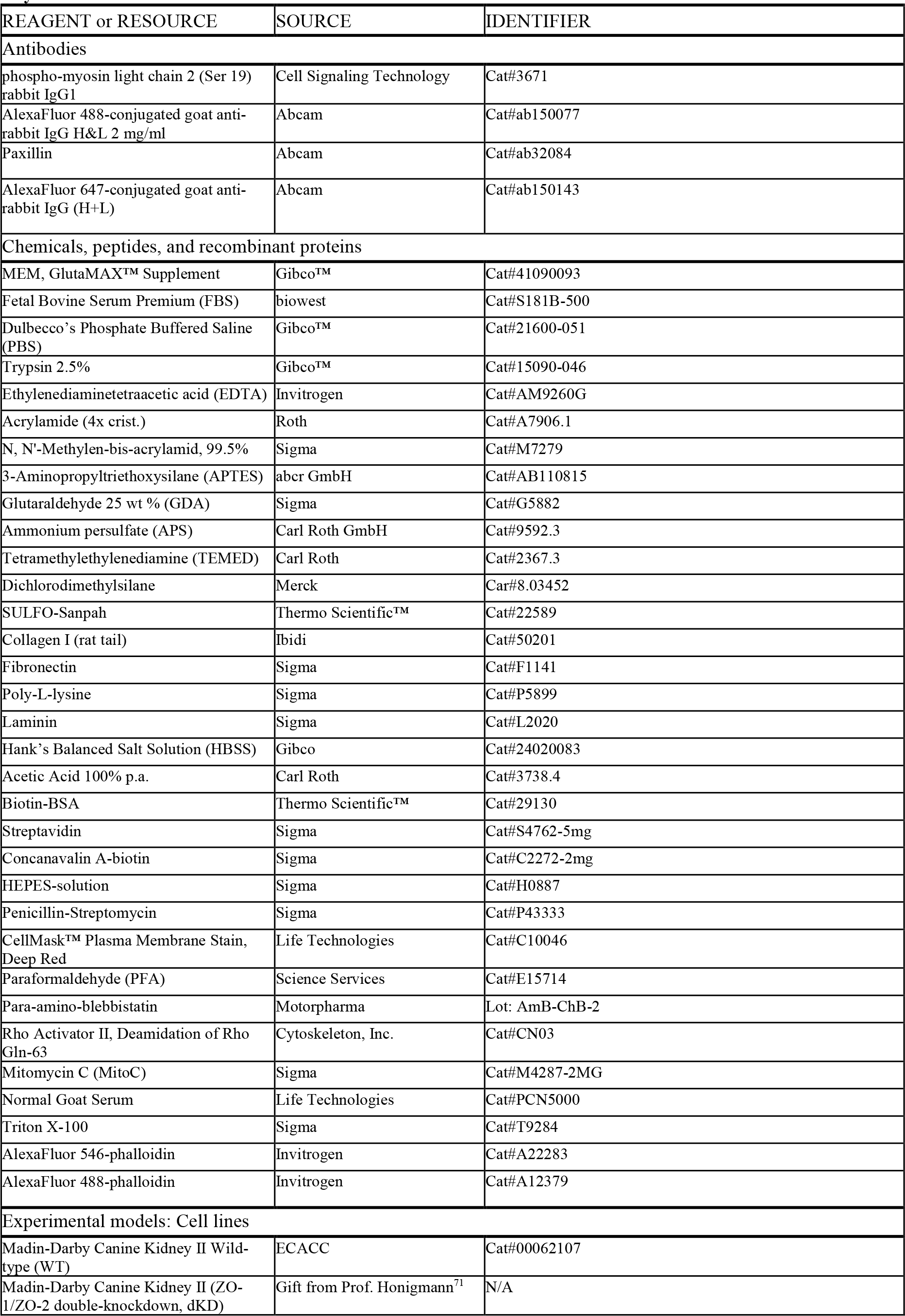
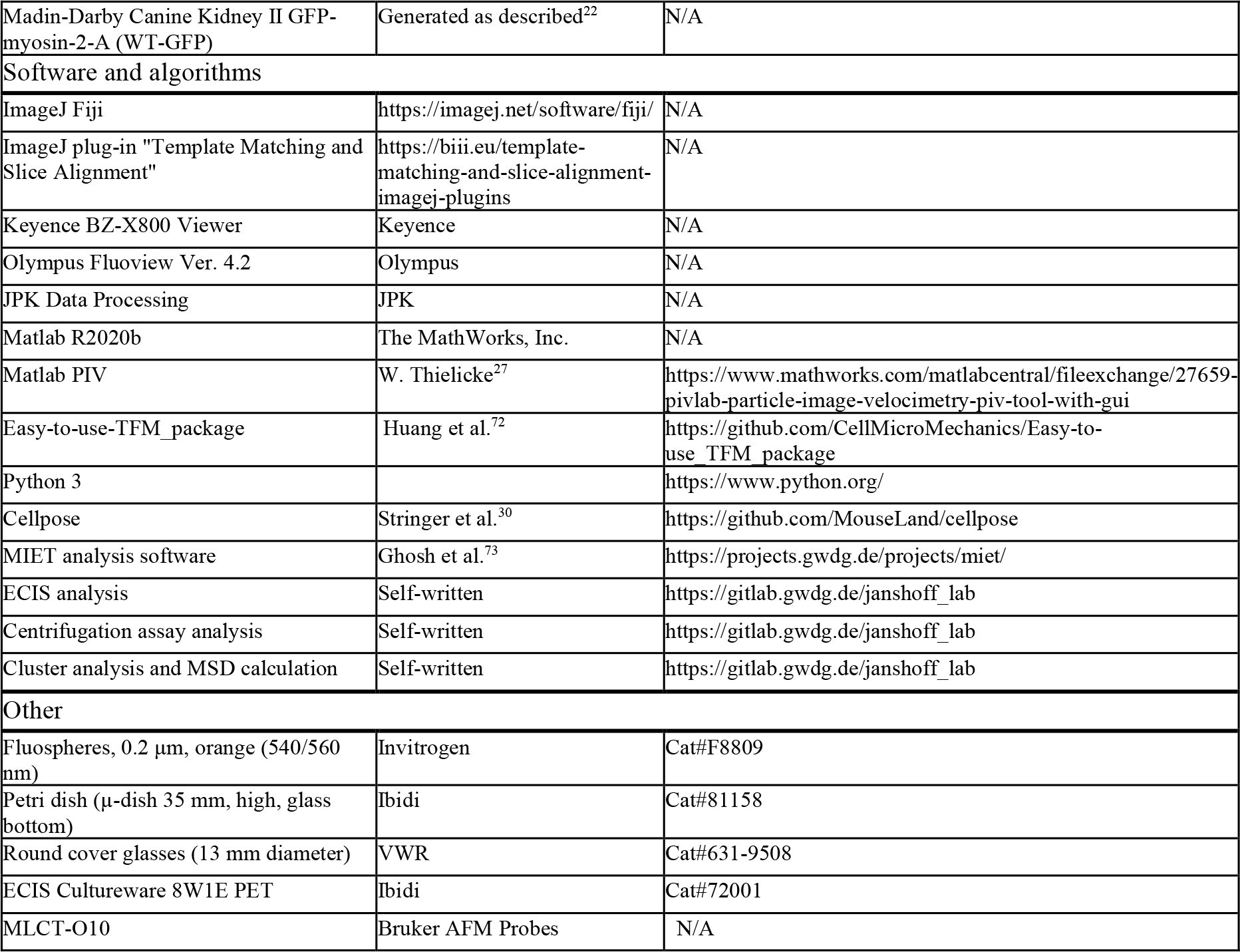

## RESOURCE AVAILABILITY

### Lead contact

Further information and requests for resources and reagents should be directed to and will be fulfilled by the lead contact, Andreas Janshoff.

### Materials availability

This study did not generate new unique reagents.

### Data and code availability

All data reported in this paper will be shared by the lead contact upon request.

Any additional information required to reanalyze the data reported in this paper is available from the lead contact upon request.

## METHOD DETAILS

### Cell culture

Madin-Darby Canine Kidney cells (strain II, MDCKII) were cultured in M10F^−^ (Minimum Essential Medium (MEM) containing Earle’s salts, 10% Fetal Bovine Serum Premium (FBS), 2 mM glutaMAX at 37 °C and 5% CO_2_ in a humidified incubator. Cells were passaged every two to three days using phosphate buffered saline pH 7.4 (PBS^−^) containing trypsin/EDTA (0.25%/0.02% w/v). The knockdown of ZO-1 and ZO-2 in MDCKII cells was done by Beutel et al. using CRISPR/Cas9.^71^ MDCKII cells expressing GFP-myosin-2-A (WT-GFP) were generated as previously described by Skamrahl et al.^22^

### Sample preparation and analysis

#### PAA gel preparation

A monomer stock solution (7.5% acrylamide (AA) and 0.075% N,N’-methylenebisacrylamide (Bis-AA)) was mixed in distilled water to prepare polyacrylamide (PAA) gels yielding a Young’s modulus of 1.61 kPa. The bottom of a Petri dish (μ-dish, 35 mm, high, glass bottom) was incubated with 150 μL of 3-aminopropyltriethoxysilane (APTES) for 5 minutes and then rinsed three times with distilled water. Subsequently, the dried surface was treated with 200 μL 2.5% glutaraldehyde solution (GDA) in PBS^−^ for 30 minutes, followed by two rinses with distilled water. Coverslips (13 mm, borosilicate glass) were washed twice with isopropanol and then cleaned thoroughly using a plasma cleaner (Zepto, Diener Electronics, Ebhausen, Germany). They were then coated with dichlorodimethylsilane for 5 minutes. Finally, the coverslips were washed with 70% ethanol and distilled water and dried completely using nitrogen gas.

A solution of 4 μL fluorescent beads (Fluospheres, 0.2 μm, orange (540/560 nm) and 246 μL of the monomer stock solution was homogenized using a sonicator for 30 min. Then, 2.5 μL ammonium persulfate (APS, 1 mg/10 μL) and 0.5 μL tetramethylethylenediamine (TEMED) were added to the solution and mixed thoroughly. 6 μL of the solution was added to each prepared Petri dish, covered with a coverslip, and allowed to polymerize for 45 min in a humid atmosphere. The coverslip was carefully removed after the gel was fully polymerized. 200 μL SULFO-Sanpah solution (0.5 mg/mL) was added to the gels and exposed to UV light (365 nm) for 8 min. Subsequently, the gels were rinsed with distilled water. These steps were repeated twice. Afterward, the gels were coated with collagen (1 μL of collagen I (rat tail in PBS^−^), 5 mg/mL in 199 μL PBS^−^ with 0.02 M acetic acid) for one hour at room temperature. The gels were stored for further use in distilled water at 4°C.

#### Migration assay

For the migration experiments, Petri dishes containing PAA gel without fluorescent beads were used onto which 5×10^5^ cells (for co-culture, a 1:1 mixture of WT/dKD was used) were seeded in 300 µL M10F^−^, to ensure the cells only attach to the gel. After 2 hours, the cells were rinsed with M10F^−^ before 2 mL was added for incubation and imaging. Three to four hours after seeding, the cells were placed in the incubation system (Uno-T-H-CO2; Okolab S.R.L., Pozzuoli NA, Italy) mounted on an inverted optical microscope (BZ-X810; Keyence, Neu-Isenburg, Germany) equipped with a 20× phase-contrast objective (Plan Fluor ELWD 20x; Nikon Europe B.V., Amstelveen, The Netherlands). The incubation chamber was operated at 37 °C and 5 % CO_2_ with sufficient humidity. Phase contrast images were acquired every 7.5 min over a period of 48 h. Focus tracking was applied, and three vertical images were selected in a range of 8 μm to account for vertical drift effects, of which the best-focused images were selected manually for analysis. The phase contrast videos obtained were analyzed using particle image velocimetry (PIV) with the MATLAB application PIVlab to compute the velocity vectors of each frame.^27^ The velocity vectors were generated with an interrogation window of 32×32 pixels (24.64×24.64 µm^2^), and the average velocity was defined as the mean velocity vectors in one frame. Mean and standard deviation were determined from independent experiments: WT (*m* = 27, *n* = 9), dKD (*m* = 22, *n* = 6), WT + Blebb (*m* = 15, *n* = 4), dKD + Blebb (*m* = 12, *n* = 4), WT + CN03 (*m* = 6, *n* = 1), dKD + CN03 (*m* = 6, *n* = 1).

#### Cell segmentation

The Cellpose cell segmentation algorithm was used to generate a mask for each cell.^30^ To accurately capture all cells in the layer, parameters for flow and cell probability were set to 1 and −6, respectively. Each cell’s diameter (short and long axis) and the cell density in one frame were then calculated from the cell masks obtained from each frame.

#### Eccentricity analysis

The eccentricity (defined as the ratio of the length of the short (minor) axis to the length of the long (major) axis) was calculated from the Cellpose data and averaged for each frame. Mean and standard deviation were determined from independent experiments: WT (*m* = 18, *n* = 5), dKD (*m* = 15, *n* = 4), WT + CN03 (*m* = 6, *n* = 1), dKD + CN03 (*m* = 6, *n* = 1).

#### Cluster Analysis and MSD

For cluster analysis and mean squared displacement (MSD) calculations, we utilized an optical flow analysis tool that we developed in Python. This tool is based on an algorithm originally proposed by Gunnar Farnebäck.^74^ Optical flow analysis generally uses the changes in the brightness pattern in a time series of images to determine the distribution of apparent velocities. The algorithm is tuned using two Gaussian kernel parameters, one for blurring the images in the polynomial fitting step (blurring kernel) and one for the flow field’s smoothness in the algorithm’s weighted neighborhood estimation (flow field kernel). To identify the best-suited parameters, an example image from the video stack is rotated by a defined angle, causing a well-defined displacement field. Then, the difference between the analytical displacement field and the calculated result from the optical flow algorithm is determined for multiple sets of parameters. A set of parameters is found reasonable when the standard deviation between the analytical and calculated fields is of the order of 0.3 px. Typical parameters used for these experiments were *σ* = 4 for the blurring kernel and *σ*_flow_ = 2 for the flow field kernel. The images are sharpened before optical flow calculation by adding the difference between the original and a Gaussian blurred version created by a convolution of a five-pixel Gaussian kernel.

Subsequently, a tracking approach is implemented to recreate example trajectories from the detected changes in the brightness patterns. First, a tube filter, using the eigenvalues of the Hessian matrix, extracts points in the image that are part of the high-contrast cell borders. This ensures that only pixels belonging to cell borders are tracked, ignoring noise from cell interiors. Randomly chosen points from the cell borders of the first frame served as starting points. Using the optical velocity fields, the following location of each chosen starting pixel is added to the trajectory. The algorithm iterates through the velocity fields and adds the velocities until the accumulated displacement is sufficient to assign the point to a neighboring pixel.

This way, the locations and velocities along a pixel path are gathered at subpixel accuracy. The list of velocities along the trajectories along a pixel path are used as displacements for calculating mean squared displacement (MSD), as they are not restricted pixel integer values. For cluster analysis, the estimated Cellpose (description see above) labels are assigned their average optical flow angle. The gradient is calculated from this image, corresponding to zero inside each label and the angular difference at the border. The borders with less considerable differences than 10° are removed. The remaining regions correspond to ensembles of cells moving together. Mean and standard deviation were determined from independent experiments: WT (*m* = 20, *n* = 7), dKD (*m* = 17, *n* = 5)).

#### Electric cell-substrate impedance sensing

The monolayer’s impedance was measured with an ECIS setup ZΦ (Applied Biophysics, NY, USA). The measurement system consists of an 8-well cell culture dish with gold electrodes deposited on the bottom of each well (8W1E array) attached to a lock-in amplifier with an internal oscillator and computer controlled relays to switch between the different wells. Each well of the 8-well electrode array has a small circular working electrode (area: 5×10^−4^ cm^2^) and a large counter electrode (area: ∼ 0.15 cm^2^) in coplanar geometry. The small electrode is formed by a circular opening (Ø = 250 μm) in a photoresist layer that insulates the rest of the gold film from the bulk electrolyte. The ultrasmall electrode ensures that the system’s total impedance is solely attributed to the cells grown on the small working electrode, thereby maximizing the sensitivity to detect subtle morphological alterations in the cell layer. Impedance data are obtained by applying an AC signal at a given frequency with 1 V amplitude through a 1 MΩ shunt resistor using the internal oscillator, leading to an approximate current of merely 1 µA running through the cell layer depending on the frequency. The resulting in-phase and out-of-phase voltages across the system are measured by a phase-sensitive lock-in amplifier and converted to the complex impedance of the system.

After adding 200 µL M10F^−^ to each chamber of an 8W1E array, the array was kept in a humidified incubation chamber (37 °C, 5% CO_2_). Cells were passaged as described before and seeded into each chamber (2×10^5^ cells in 200 µL M10F^−^). The impedance of cell-covered electrodes was measured using the multiple frequencies over time (MFT) mode at eleven frequencies ranging from 62.5 Hz to 64 kHz with a temporal resolution of approximately 2.6 minutes.

For the second data set (SI Figures 2B, 2C, and 4), 2×10^5^ (low cell density) and 4×10^5^ (high density) were seeded in 400 µL M10F^−^ per chamber. The electrodes were immersed with collagen I rattail (5 µg/cm^2^ dissolved in PBS^−^ with 0.02 M acetic acid), fibronectin (3 µg/cm^2^ dissolved in Hank’s Balanced Salt Solution (HBSS)), laminin (2 µg/cm^2^ dissolved in HBSS) or poly-L-lysine (4 µg/cm^2^ dissolved in distilled water) for 60 minutes at room temperature. The electrodes were then thoroughly rinsed three times with PBS^−^. Mean and standard deviation were determined from independent experiments: WT low density (*m* = 11, *n* = 2), WT high density (*m* = 11, *n* = 2), dKD low density (*m* = 13, *n* = 2), dKD high density (*m* = 11, *n* = 2), WT + Blebb (*m* = 10, *n* = 2), dKD + Blebb (*m* = 10, *n* = 2), WT + CN03 (*m* = 14, *n* = 2), dKD + CN03 (*m* = 14, *n* = 2).

#### Cell-substrate distance fluctuation analysis

For recording the cell-substrate distance fluctuations of adherent cells reflecting shape fluctuations comprising predominately cell-substrate distance variation, the complex resistance at discrete frequencies was recorded.^75^ At 4 kHz the relative impedances of both cell lines match and are governed by shape fluctuations of the cells. Only confluent layers were considered to rule out cell-number fluctuations on the electrode. A period of 25 hours was selected for the analysis. These trimmed data sets contain both cell-substrate distance fluctuations and an overall trend due to migration and division. To identify and remove this overall trend, we used the empirical mode decomposition method. This signal decomposition yields a frequency-resolved distribution of modes in which the very low-frequency modes display the overall trend. Consequently, subtracting the low-frequency modes from the original signal obtains a detrended signal containing only cell-substrate distance fluctuations. Quantitative evaluation of cell-substrate distance fluctuations as a time series is performed by determining the variance of these detrended signals (detrended variance analysis, DVA) and detrended fluctuations analysis as outlined below.

#### Detrended Fluctuation Analysis

DFA analysis, as outlined by Peng et al.,^39^ involves the integration and partitioning of datasets of length *N* into *l* boxes with a box size of *n* within each box, a linear trend was subtracted, and the variance *F*(n) was computed by evaluating the difference between the original data *y*(k) and the detrended data *u*(k) using:

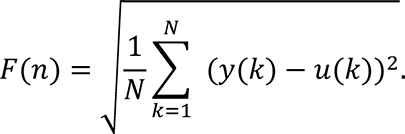

The relationship between *F*(n) and *n* was then visualized in a ln(*F*(n)) - ln(n) plot. The slope of the linear fit applied to this plot provides direct insights into the correlation of the analyzed system. α_DFA_ values and the associated Hurst coefficient *H*, related to α_DFA_ (α = *H* + 1 for fractional Brownian motion processes with long term memory), indicate temporal persistence in time series. While for short-range processes, the coupling between successive values decays quickly with distance in an exponential fashion, long-range processes show a power-like decay of the autocorrelation. Mean and standard deviation were determined from independent experiments: WT (*m* = 16, *n* = 2), dKD (*m* = 24, *n* = 3).

#### Traction Force Microscopy

For traction force experiments, Petri dishes with beads containing PAA gels were used. 8×10^4^ cells were seeded on the gel in 300 µL M10F^−^. After 2 hours of pre-incubation in a humidified incubator (37 °C, 5% CO_2_), 2 mL M10F^−^ were added. For WT-GFP/dKD co-culture measurements, 4×10^4^ cells of each type were mixed and seeded simultaneously.

Images were acquired using a confocal laser scanning microscope (Fluoview 1200; Olympus, Japan) with a 60x oil immersion objective (UPLFLN 60XOIPH; Olympus, Japan) and a stage top incubation system (blue line; ibidi, Martinsried, Germany) set to 37 °C, 5% CO_2_. A phase contrast image of the cells (to determine cell contours using Cellpose) and a fluorescent image of the image plane containing the fluorescent beads were acquired. To remove the cells, a solution of trypsin (2.5%) with 0.05 M EDTA in PBS^−^ was incubated for 10 to 20 minutes until the cells had completely detached before a second image of the bead-containing plane was acquired. The ImageJ plug-in “Template Matching and Slice Alignment” was used to remove sample shifts during the measurement. The MATLAB application PIVlab was used to calculate the displacement field of the beads.^27^ The displacement was determined using four query windows (256 px, 128 px, 64 px, 32 px), and the forces were then calculated from the displacement fields.^72^ A mesh size of 16 pixel, a Poisson’s ratio of 0.48, and integrated manual noise selection were used. Each gray dot in the plots indicates an independent experiment.

#### Cell-substrate adhesion measurements

A CellHesion 200 AFM (JPK Instruments; Berlin, Germany) was used to measure cell-substrate adhesion using a tipless cantilever (MLCT-O10, Bruker AFM Probes). To prepare the cantilever, it was first cleaned in acetone for 5 minutes and then exposed to UV light (365 nm) for 15 minutes. This was followed by overnight incubation of the cantilever (at least 12 hours) in 50 µL biotin-BSA solution (0.5 mg/mL in 0.1 M NaHCO3, pH 8.6) at 37 °C. Subsequently, the cantilevers were washed three times with PBS^−^ and incubated in 50 µL streptavidin solution (0.5 mg/mL in 0.01 M PBS^−^, pH 7.3) for 20 minutes at room temperature. After 3x PBS^−^ -wash, a 20 min incubation in concanavalin A-biotin solution (0.2 mg/mL in PBS^−^) was performed to attach cells firmly to the cantilever. The whole procedure was adopted from Wojcikiewicz et al.^76^ 6×10^4^ cells suspended in M10F^−^ medium (containing HEPES and penicillin-streptomycin) were added to a PAA gel for adhesion measurements. To attach a cell to the cantilever, a pre-adhered cell was picked with a setpoint of 2 nN and allowed to adhere to the cantilever for 15 minutes without contact with the PAA gel. Subsequently, the cell was pressed onto the substrate with a setpoint of 2 nN for 150 seconds (typical adhesion times for single cell force spectroscopy, that allow for initial adhesion but avoid apoptosis due to prolonged pressure on the cell). The approach and retract velocity were set to 0.5 μm s^−1^. The maximal adhesion force was determined using JPK Data Processing software after linear baseline correction and detection of the contact time point. Mean and standard deviation were determined from independent experiments: WT (*m* = 25, *n* = 6), dKD (*m* = 34, *n* = 6), WT + Blebb (*m* = 70, *n* = 4), dKD + Blebb (*m* = 45, *n* = 4).

#### Centrifugation assay

The centrifugation assay was performed as described elsewhere.^45,46^ 1×10^5^ Cells were seeded on top of a coverslip (13 mm diameter), which was placed in a self-built holder. The sample was placed in a centrifuge (Megafuge 16R Centrifuge, Thermo Electron LED GmbH, Langenselbold, Germany) for 5min after an incubation time of 10, 20, and 40 min with an angular velocity of 5000 rotations/min. At incubation times much longer than 40 min, cell adhesion will exceed cohesion, making removal of the cells from the substrate with the forces available here impossible. The whole area of the coverslip was imaged before and after the centrifugation with an inverted optical microscope (BZ-X810; Keyence, Neu-Isenburg, Germany) equipped with a 10× phase-contrast objective (Plan Fluor 10x; Nikon Europe B.V., Amstelveen, The Netherlands). Mean and standard deviation were determined from independent experiments: WT (*m* = 3, *n* = 3), dKD (*m* = 3, *n* = 3).

#### Metal-induced energy transfer

All MIET measurements were carried out with a custom-built confocal microscope equipped with a multichannel picosecond event timer (HydraHarp 400, PicoQuant GmbH; Berlin, Germany), allowing for fluorescence lifetime imaging. The system was equipped with a high numerical aperture objective (Apo N, 60×/1.49 NA oil immersion, Olympus) for both focusing excitation light and collecting fluorescence light. A white-light laser system (SC400-4-20, Fianium) with an acoustic-optical tunable filter (AOTFnC-400.650-TN, AA Optic) served as the excitation source (λ_exc_ = 645 nm). The excitation light was reflected by a non-polarizing beam splitter towards the objective. Back-scattered excitation light was blocked with a long-pass filter (BLP01-635R, Semrock). Collected fluorescence was focused onto the active area of an avalanche photodiode (PDM series, MPD). The fluorescence decay curves were fitted with a multi-exponential decay model (up to 10 mono-exponential components), from which the average excited state lifetime was calculated. A semitransparent metallic film consists of a 15 nm gold film deposited on a 2 nm titanium film for better adhesion to glass. The metal films were prepared by vapor deposition onto a glass bottom Petri dish (µ-Dish, Ibidi) using an electron beam source (Univex 400, Leybold, Cologne, Germany) under high-vacuum conditions (∼10^−6^ mbar). During vapor deposition, the film thickness was monitored using an oscillating quartz unit and verified using a profilometer. 5×10^5^ cells were seeded on the inner surface of the petri dish, as described above. After approximately 24 hours in the incubator (37 °C and 5% CO_2_), the cells were washed three times with M10F^−^. A solution of 1 μL CellMask Plasma Membrane Stain Deep Red with 499 μL M10F^−^ was added to the cells. After ten minutes of incubation (37 °C and 5% CO_2_), cells were washed three times with M10F^−^ and fixed in a 4% paraformaldehyde solution for 15 min. Cells were stored in PBS^−^ for measurement. The analysis of the raw data was performed with a self-written Matlab script described in Karedla et al.^73,77^ Cell center elevations were quantified by manually cropping the elevation profiles and determining a mean value. Mean and standard deviation were determined from independent experiments: WT (*m* = 6, *n* = 2), dKD (*m* = 8, *n* = 2), WT + Blebb (*m* = 6, *n* = 1), dKD + Blebb (*m* = 5, *n* = 1), WT + CN03 (*m* = 6, *n* = 1), dKD + CN03 (*m* = 7, *n* = 1).

#### Drug Treatment

Para-amino-blebbistatin (Blebb) was dissolved at 50 mM in DMSO, and the aliquots were stored at −20 °C. Cell samples were washed twice with M10F^−^ and incubated with M10F^−^ containing 50 μM of Blebb at 37 °C and 5% CO_2_ for 30 min. Afterward, three washes with M10F^−^ were performed, and the samples were prepared for the respective experiments.

A stock solution of Rho Activator II (CN03) was prepared at 0.1g/L in water and stored at - 70 °C. After washing once with M10F^−^, the cells were incubated with 1 μg/mL CN03 (in MEM) at 37 °C and 5% CO_2_ for 3.5 h. Subsequently, the samples were washed three times with M10F^−^ and prepared for further experiments.

A stock solution of Mitomycin C (MitoC) was prepared in water (500 μg/mL) and stored in aliquots of 150 μL at −20 °C. Samples were seeded as before and treated with 10 μg/mL MitoC in an incubator at 37 °C, 5% CO_2_ for 1 hour. The samples were washed three times with M10F^−^ and used for subsequent experiments.

#### Staining (actin, myosin)

Cells were seeded on the PAA gels as described above. After 24h, samples were washed three times with PBS and then incubated for 15 min in PFA (4% (w/v) in PBS^−^) to fixate cells. Then, permeabilization was performed with Triton X-100 (0.1% (v/v) in PBS^−^) for 15 minutes. After three subsequent washes with PBS^−^ blocking/dilution buffer (PBS^−^ containing 2% (v/v) Normal Goat Serum) was added for 30 minutes.

Samples were incubated with a primary antibody (Phospho-myosin: 2 μg mL-1 (1:50 (v/v)) light chain 2 (Ser 19) rabbit IgG1) diluted in blocking/dilution buffer for 1 h. After incubation with the primary antibody, the samples were briefly rinsed with PBS^−^ before washing three times with PBS^−^ (each 5min on a shaker plate). Then, the secondary labeling solution (in blocking/dilution buffer) with the second antibody (1:1000 (v/v) AlexaFluor 488-conjugated goat anti-rabbit IgG H&L) and AlexaFluor 546-phalloidin (165 nM) was incubated with the sample for 1 h. Samples were washed three times with PBS^−^ (each 5min on a shaker plate) and kept in PBS^−^ for imaging.

#### Staining (paxillin)

To fix the cells, samples were rinsed once PBS^−^ and then fixed with 4% PFA for 20 min. After fixation, cells were permeabilized with 0.1% Triton solution for 15 min. After rinsing three times with PBS^−^, samples were incubated in blocking/dilution buffer for 60 min to block unspecific binding sites. Samples were incubated in primary antibody (anti-paxillin) diluted in blocking/dilution buffer overnight at 4 °C (1:200 (v/v)). Subsequently, cells were washed three times with PBS^−^ on a platform shaker (5 min). The secondary antibody (AlexaFluor 647-conjugated goat anti-rabbit IgG) was diluted with blocking/dilution buffer (1:1000 (v/v)) and incubated together with AlexaFluor 488-phalloidin (1:40 (v/v)) for 2 h. Following the secondary antibody, samples were washed three times with PBS^−^ (5 min) on a shaker and kept in PBS^−^ for imaging.

#### Actin and myosin intensity quantification

Images of the basal side were taken on the confocal laser scanning microscope (Fluoview 1200, Olympus, Japan). Myosin staining was imaged with a 60x oil immersion objective (UPLFLN 60XOIPH, NA1.25, Olympus, Japan) and actin staining with a 100x oil immersion objective (UPLXAPO 100x/NA1.45, Olympus). Data sets were created on the same day with the same settings. Myosin intensity was measured in the cell center. For this, the corresponding phase contrast images were segmented by Cellpose to create a mask of the cell borders. ImageJ was subsequently used to analyze the corresponding intensities. Actin intensity measurements were analyzed by ImageJ averaged over an entire image. Mean and standard deviation were determined from independent experiments: WT myosin (*m* = 4, *n* = 1), dKD myosin (*m* = 3, *n* = 1), WT actin (*m* = 17, *n* = 2), dKD actin (*m* = 16, *n* = 2), WT actin + CN03 (*m* = 16, *n* = 1), dKD actin + CN03 (*m* = 16, *n* = 1).

#### Quantification of focal adhesions

Images of the basal side were taken on the confocal laser scanning microscope (Fluoview 1200, Olympus, Japan). Focal adhesions were imaged with a 60x oil immersion objective (UPLFLN 60XOIPH, NA1.25, Olympus, Japan). Data sets were created on the same day with the same settings. Binary images of paxillin were created by thresholding confocal pictures. White spots indicate focal adhesions. The number of white spots and their average size were determined for each picture. Mean and standard deviation were obtained from independent experiments: WT (*m* = 7, *n* = 1), dKD (*m* = 14, *n* = 1).

#### Quantification and statistical analysis

All data were written in the text as mean ± standard deviation. The gray dots in the box plots represent independent experiments (biological and technical replicates). Box plots show the median with the 25th and 75th percentiles, and whiskers show the 5^th^ and 95^th^ percentiles. A significance test (Wilcoxon rank-sum test) was performed for all data sets, denoted by the *p*-value. A *p*-value of < 0.1 was considered significant and denoted by one asterisk (*). *p* < 0.05 and *p* < 0.005 were indicated by two (**) and three (***) asterisks, respectively. The number of biological replicates (*n*) and technical replicates (*m*) of the experiments is given in the respective method sections.

