## Supplemental Information for "Cell-substrate distance fluctuations of confluent cells enable fast and coherent collective migration"

### Supplementary information

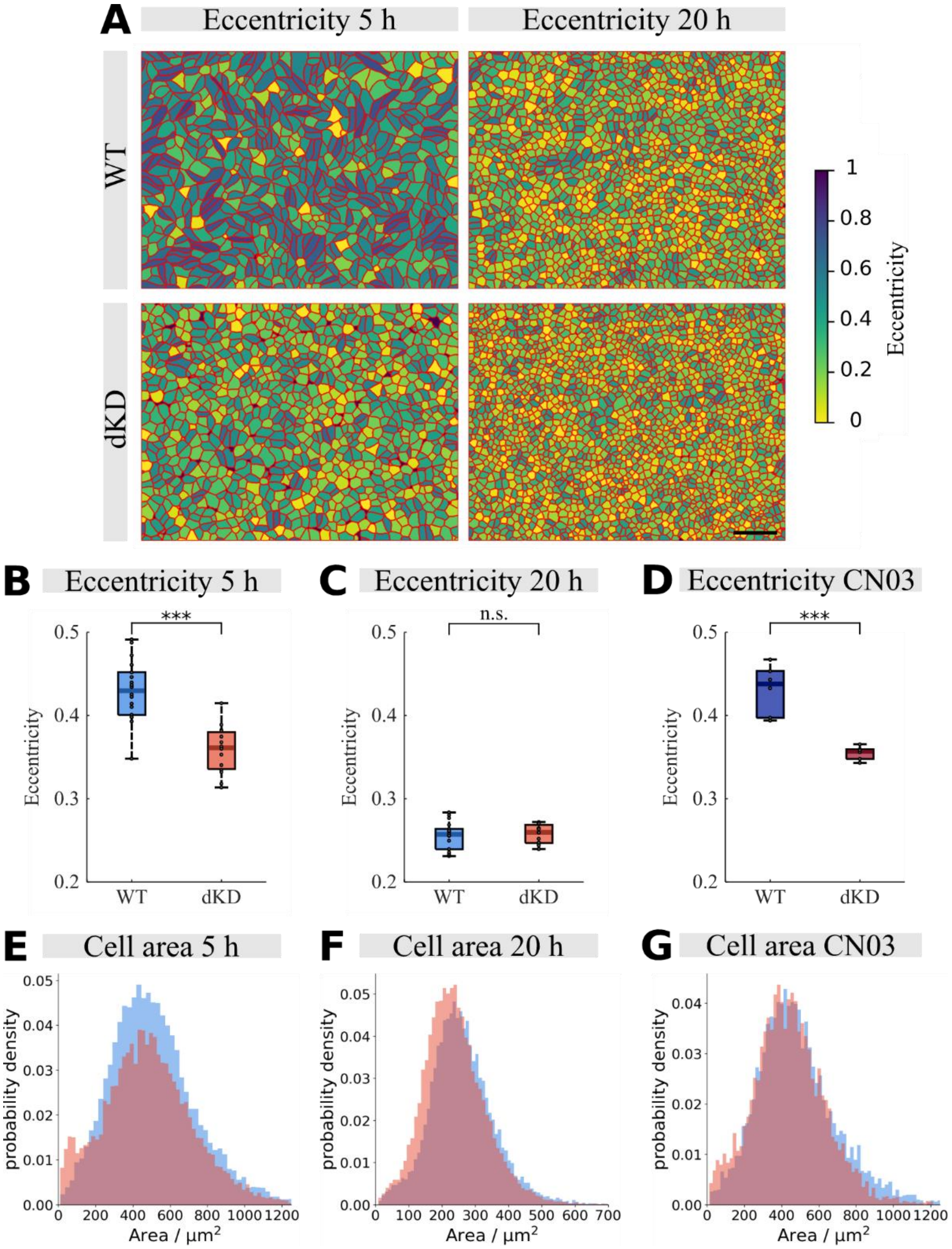


**SI-Figure 1. Cell eccentricity analysis and cell size over time.** A) Representative eccentricity maps of WT and dKD monolayers. Cell outlines obtained from Cellpose cell segmentation are overlayed in red. Scale bar: 150 µm. B) Eccentricity of WT and dKD monolayers five hours after seeding. C) Eccentricity of WT and dKD monolayers 20 hours after seeding. D) Eccentricity of CN03 treated WT and dKD monolayers five hours after seeding. E) Cell sizes of WT and dKD monolayers five hours after seeding. F) Cell sizes of WT and dKD monolayers 20 hours after seeding. G) Cell sizes of CN03 treated WT and dKD monolayers five hours after seeding.

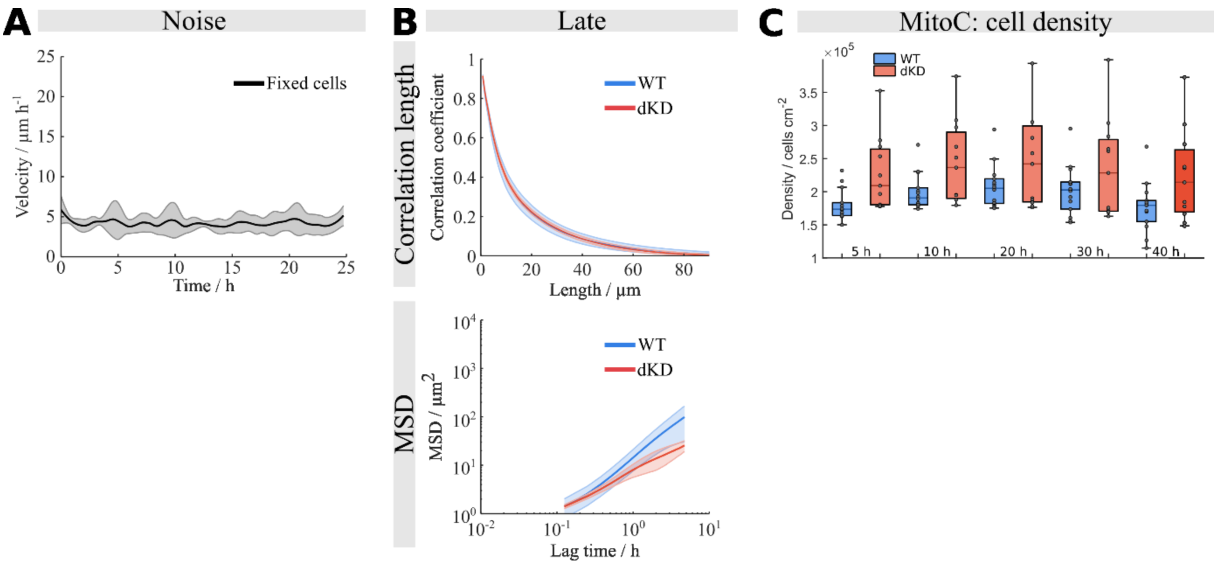


**SI-Figure 2.** A) Averaged time-resolved velocity of a fixes WT sample showing the "baseline velocity". B) Upper plot: spatial correlation of velocity for late (jammed) state. Lower plot: Mean square displacements (MSD) of WT and dKD at late (jammed) state. C) Cell densities of Mitomycin C treated WT and dKD monolayers over time.

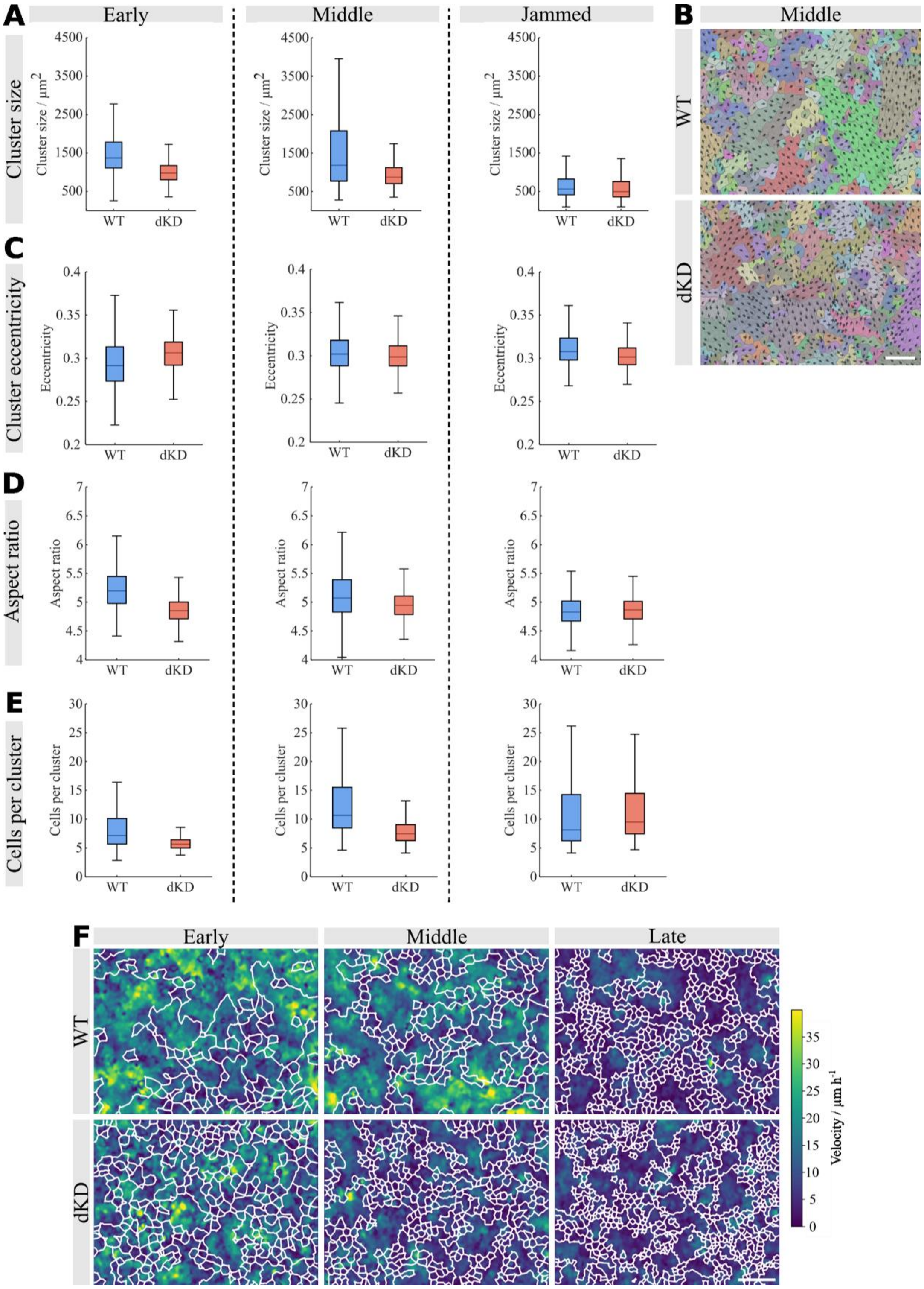


**SI Figure 3. Analysis of cluster sizes.** A) Size of cell clusters that are composed of cells with the same preferred orientation of velocity vectors of early, middle and jammed states. B) Representative motion cluster maps of WT and dKD with corresponding cell outlines obtained by Cellpose at middle (between 15 h and 20 h after seeding) phases of the observation period. The arrows represent the motion of individual cells. Cells with similar directionality are grouped into a cluster with the same color. Scale bar: 150 µm. C) Cluster eccentricities of early, middle and jammed states. D) The aspect ratio of the clusters of early, middle and jammed states. E) The average number of cells per cluster of early, middle and jammed states. F) Representative velocity maps generated from the corresponding velocity vectors obtained from PIV with cell cluster outlines (white) obtained from the cell cluster analysis. WT cells form larger clusters that are clearly faster. Scale bar: 150 µm.

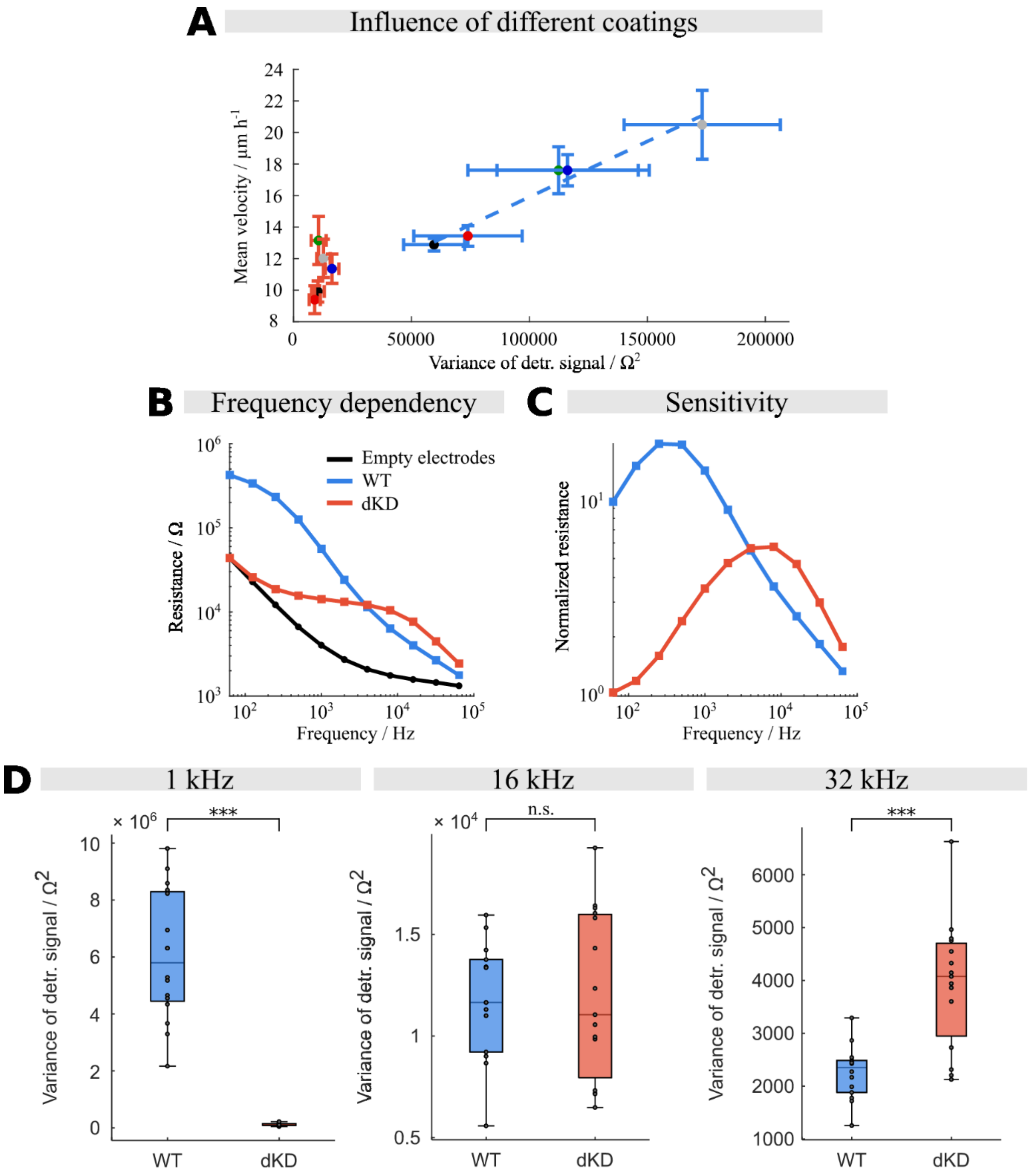


**SI-Figure 4. Frequency dependency and influence of different coatings.** A) The average speed of WT and dKD cells is compared to the respective variance in the detrended signal from the cell layer post wound closure. Each point indicates a different substrate coating: black: no coating; green: collagen; gray: fibronectin; blue: poly-L-lysine; red: laminin. A linear regression of the WT samples is shown with the dashed line ($R$ = 0.978). B) Mean impedances (the complex impedance is divided into a real part (resistance) and an imaginary part (capacitance) in series) of WT, dKD and empty electrodes as a function of frequency. C) Resistance normalized to that of the empty electrode to identify the maximal response of the cells on the electrode. The cell-substrate-fluctuations were evaluated at 4 kHz, where both curves intersect. D) Detrended variance analysis of cell-substrate fluctuations measured with ECIS at lower and higher frequencies illustrating that in the relevant low frequency regime WT cells display larger fluctuations.

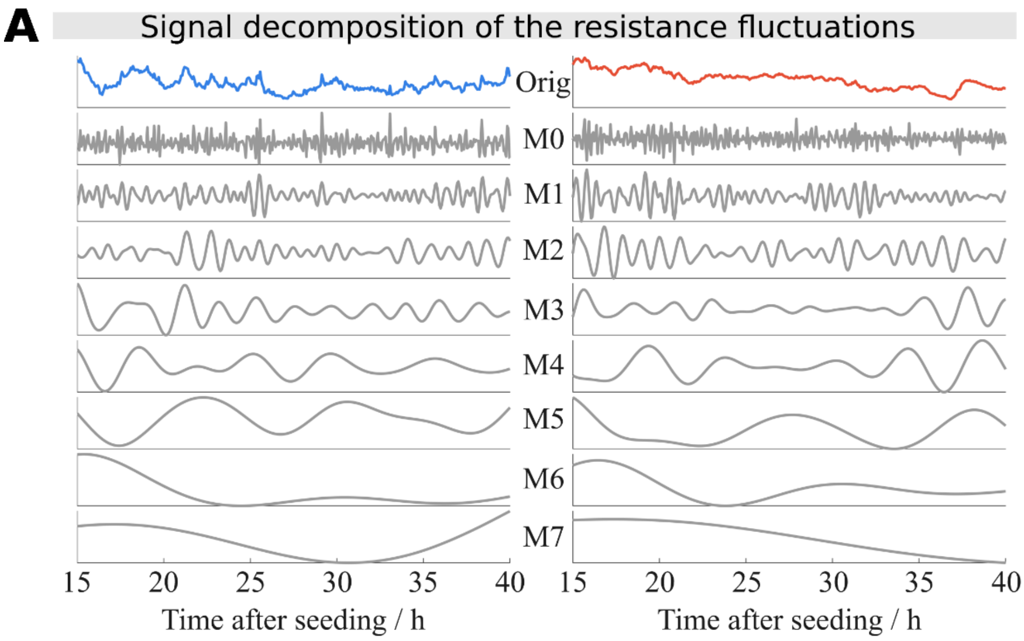


**SI Figure 5. Signal decomposition of the resistance fluctuations.** A) Signal decomposition of the resistance fluctuations in representative WT and dKD monolayer samples (Orig). The signal decomposition ranges from high (M0) to low (M7) frequencies.

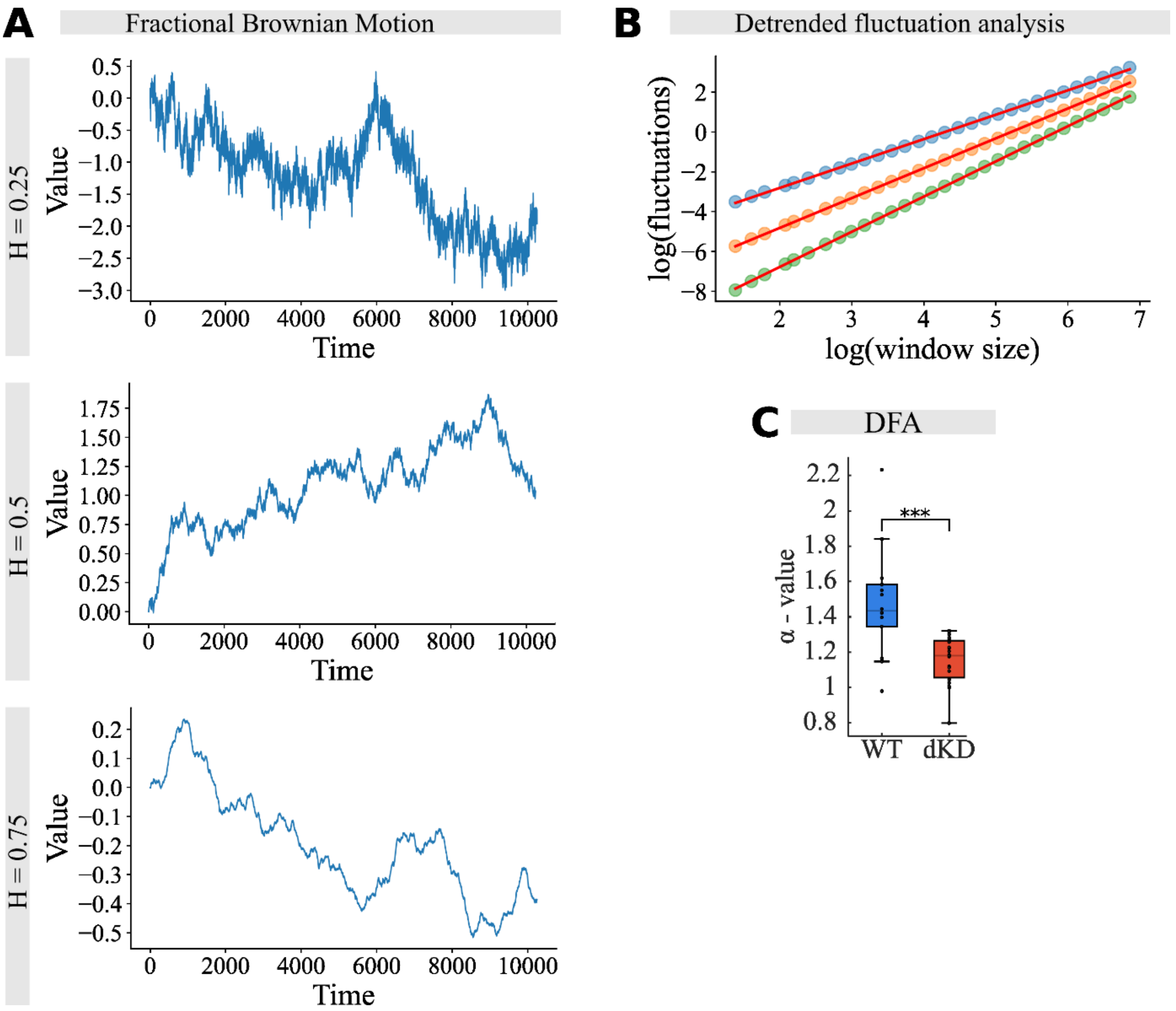


**SI-Figure 6. DFA analysis.** A) Simulated time series of fractional Brownian motion with preset Hurst coefficients ($H$ = 0.5 refers to Brownian motion). B) DFA-plots of the time series shown in A) showing that with increasing Hurst coefficient the slope increases (blue: $H$ = 0.25, orange: $H$ = 0.5, green, $H$ = 0.75). Higher slopes and larger Hurst coefficients ($\alpha = H+1$) indicate a longer memory. C) DFA analysis of confluent WT and dKD cells. A time trace of 10 hours after reaching confluency was chosen for the analysis.

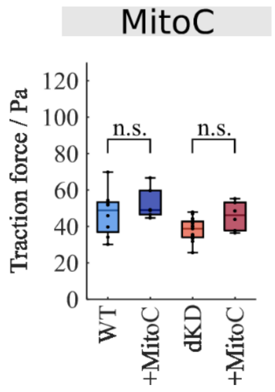


**SI-Figure 7**. Average traction forces for WT and dKD monolayers treated with Mitomycin C (MitoC) to inhibit cell division. Monolayers were treated with MitoC after 20 hours and incubated for an additional 24 hours. The cell densities of Mito C treated monolayers (WT: 3.9 ± 0.6 x $10^5$ cells/cm$^2$, dKD: 3.9 ± 0.8 x $10^5$ cells/cm$^2$) are in the same range as those observed in the untreated samples (cell density: 2.8 – 4.5 x $10^5$ cells/cm$^2$).
